# An integrative approach uncovers genes with perturbed interactions in cancers

**DOI:** 10.1101/733485

**Authors:** Shilpa Nadimpalli Kobren, Bernard Chazelle, Mona Singh

## Abstract

A major challenge in cancer genomics is to identify genes with functional roles in cancer and uncover their mechanisms of action. Here, we introduce a unified analytical framework that enables rapid integration of multiple sources of information in order to identify cancer-relevant genes by pinpointing those whose interaction or other functional sites are enriched in somatic mutations across tumors. Our accompanying method PertInInt combines knowledge about sites participating in interactions with DNA, RNA, peptides, ions or small molecules with domain, evolutionary conservation and gene-level mutation data. When applied to 10,037 tumor samples across 33 cancer types, PertInInt uncovers both known and newly predicted cancer genes, while simultaneously revealing whether interaction potential or other functionalities are disrupted. PertInInt’s analysis demonstrates that somatic mutations are frequently enriched in binding residues and domains in oncogenes and tumor suppressors, and implicates interaction perturbation as a pervasive cancer driving event.

(Software at http://github.com/Singh-Lab/PertInInt.)

## Introduction

Large-scale, concerted oncogenomic consortia have recently sequenced an unprecedented number of tumor genomes from thousands of patients across tens of cancer types [1, 2]. Analyses of these datasets promise the opportunity for improved diagnosis and additional insights into the genetic underpinnings of a staggeringly complex and heterogeneous disease [3]. More broadly, the comprehensive detection of cancer-driving mutational events, coupled with a mechanistic understanding of their functional impact, has the potential to expand our knowledge of altered cellular processes in tumors, to reveal actionable, genetic similarities between different cancer types, and to inform how evolving, heterogeneous populations of tumor cells may impact therapeutic efficacy [4–6].

A crucial first step toward these goals—differentiating the small fraction of somatic mutations with functional roles in cancer (“drivers”) from the preponderance of neutral “passenger” mutations—still poses a substantial computational obstacle [7]. While initial attempts to uncover cancer drivers at the gene level based on frequency of mutation across tumor samples have been fruitful [8, 9], such gene-centric, recurrence-based approaches are inherently unable to detect infrequently mutated driver genes and also cannot distinguish amongst mutations within the same gene that may lead to distinct tumor phenotypes or clinical responses [10]. In order to address the critical need to detect and interpret rare mutational driver events at the subgene level [11], an emerging class of approaches has begun to combine somatic mutation information with additional knowledge regarding protein site functionality, derived from analyses of evolutionary conservation [12–14], three-dimensional structure [15–20], domains [21, 22], or post-translational modification [23, 24]. These methods, however, tend to consider whether somatic mutations alter just a single type of functionality, whereas somatic mutations within putative driver genes have been found to disrupt a broad range of protein functionalities. On the other hand, machine learning approaches to classify cancer drivers incorporate multiple types of information, but due to their “black box” nature, mechanistic interpretations of their predictions are not possible [25, 26].

We and others have previously demonstrated that detecting proteins that harbor somatic mutations in their interaction interfaces is a particularly effective approach to pinpoint infrequent driver mutations as well as reason about their molecular impacts and therapeutic sensitivities [16, 17, 27–32]. Indeed, several cancer driver genes, including *TP53* and *IDH1*, are well known to harbor mutations within their interaction sites [29]. While traditionally interaction sites have been identified directly for the small fraction of human genes with actual or modeled co-complex structures, we have recently developed a domain-based approach that accurately detects residues that interact with DNA, RNA, peptides, ions or small molecules across 63% of human genes [33]. A robust computational framework that utilizes this vastly expanded knowledge-base about interaction sites and explicitly integrates it with additional lines of evidence regarding subgene functionality would provide a powerful new approach not only to detect but also to interpret a wide range of mutations driving protein dysfunction in cancer.

Here, we introduce a fast, interpretable and easily extendable framework that enables us to uncover whether somatic mutations within genes are enriched in sites associated with high measures of “functionality” as determined by multiple, possibly correlated, lines of evidence. Our implementation **PertInInt** (pronounced “pertinent,” **Pert**urbed **In Int**eractions) incorporates interaction site information, along with evolutionary conservation and domain membership information, as each of these measures informs which sites are important for protein functioning. Importantly, we derive analytical calculations that obviate the need to perform time-prohibitive permutation-based significance tests, thereby making it feasible to integrate, in a principled manner, these distinct measures of subgene-level functionality. Further, we extend our framework to consider whole-gene mutation rates, as genes that are recurrently mutated across tumors are often found to be causally implicated in cancers [34]. While other approaches have combined the output of multiple programs post hoc (e.g., [6]), PertInInt is, to the best of our knowledge, the first approach that integrates multiple alternate sources of subgene resolution data with whole gene mutational frequency within a single unifying framework in order to detect, evaluate, and infer the molecular impact of patterns of somatic mutations within all human genes.

We apply PertInInt to somatic missense mutation data arising from 10,037 tumor samples across 33 cancer types to identify genes with the most enriched mutational patterns. We find that while each source of information—interaction, domain, evolutionary conservation and whole-gene mutation frequency—is individually predictive of cancer genes, PertInInt uncovers more comprehensive sets of cancer-relevant genes when considering all sources of information together. We demonstrate that PertInInt is able to identify even those cancer genes with relatively low overall mutation rates, and that PertInInt readily outperforms previous methods while additionally revealing whether and what type of interaction potential is perturbed. PertInInt finds that numerous known oncogenes and tumor suppressors have an enrichment of somatic mutations within their interaction interfaces and, in addition, newly predicts cancer-relevant genes along with their altered interaction functionalities. Altogether, PertInInt provides a new and highly effective integrative framework to analyze large-scale cancer somatic mutation data and further our understanding of the molecular mechanisms driving cancers.

## Results

### Overview of the PertInInt framework

PertInInt aggregates somatic mutational data observed across tumor samples and identifies for each gene whether certain types of its functional sites are enriched in somatic mutations and/or whether the gene exhibits a high mutation rate across its length. We briefly overview our approach next (Fig. 1); more details can be found in the Methods section.

**Figure 1:**
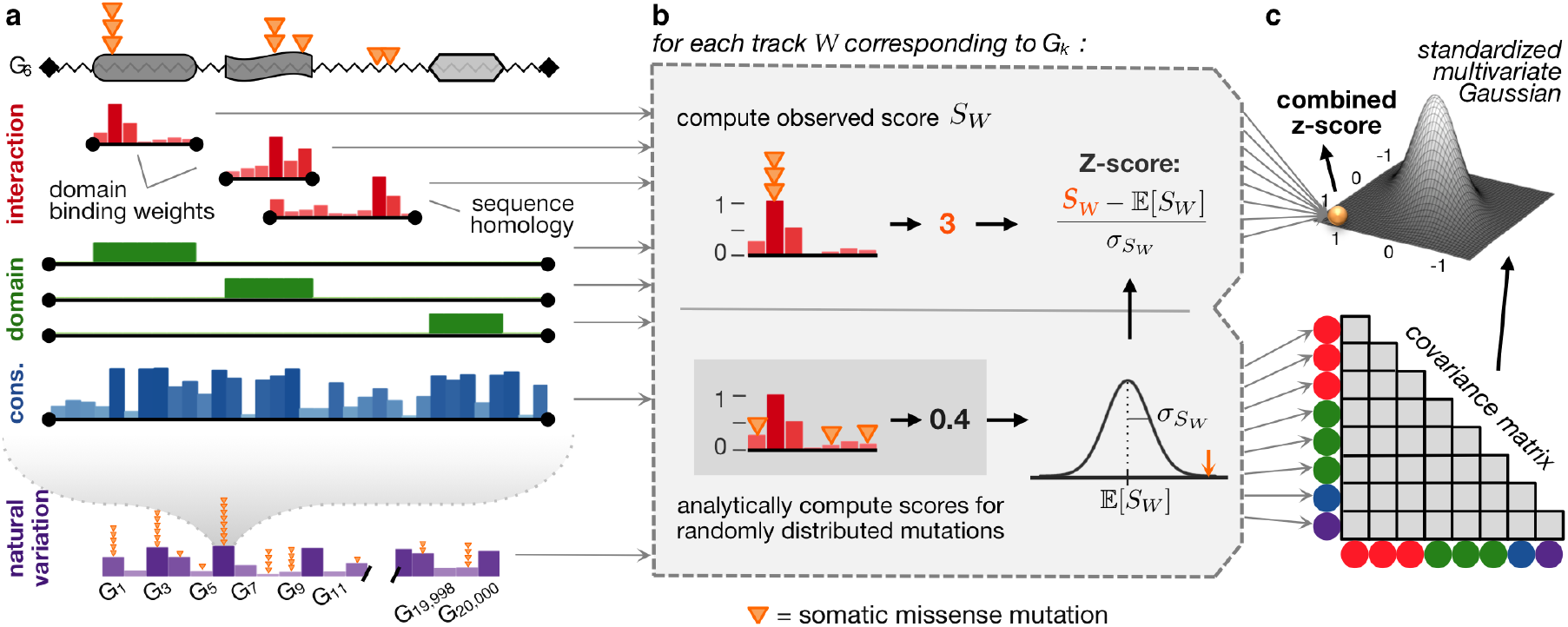
PertInInt uncovers cancer driver genes by integrating subgene per-site interaction, domain, and conservation information with whole-gene mutation frequency data. **(a)** Somatic mutations (orange triangles) affecting a protein sequence (jagged line) with three domains (grey regions) are evaluated with respect to different measures of functionality, each represented as a “track”. In interaction tracks (red), positions that are more likely to participate in ligand interactions have higher weights (vertical bars). Interaction tracks arise from domain-based binding potential calculations [33] (top two red tracks, each covering the length of the respective domain) or homology modeling [30] (bottom red track, covering the length of the modeled region). Domain tracks (green) specify which residues within a protein are part of a specific domain by 0/1 positional weights; here we have a track for each domain within the sequence. The conservation track (blue) weights each position by its evolutionary conservation across species. The natural variation track (purple) models how much each gene varies across healthy populations; here the height of the vertical bars indicates the background mutation probability rather than a per-gene weight, which is 1 for the gene being considered and 0 otherwise. **(b)** For each track *W*, we compute the score *S*_*W*_ of the observed somatic mutations as the sum of the track weights for the positions where they appear (top). To determine whether this score is higher than expected, we consider a model where somatic mutations are shuffled across the positions of the track, and the expected score (𝔼[*S*_*W*_]) and the standard deviation of the scores 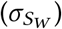 are computed and used to estimate per-track *Z*-scores (bottom); note that in our framework these values are computed analytically instead of relying on the shuffles. **(c)** *Z*-scores for all tracks are combined after analytically determining a background covariance model.

Each functional region within a protein is modeled as a “track,” and each position within a track has a corresponding 0 to 1 weight that reflects its importance with respect to the track (Fig. 1a). For each track, we compute the score of the somatic mutations with respect to the track as the sum of the weights of the positions that the mutations fall into (Fig. 1b). To determine whether the score for a track is more than expected by chance, we could shuffle the mutations across the positions of the track, and use the mean and standard deviation computed from these permutations to compute a *Z*-score; however, the mean and standard deviation for each track can be computed analytically (see Methods S3 for derivations). For each protein, we next combine the information from each of its tracks. Because tracks can overlap along the length of the protein sequence, and the somatic mutations that fall in each of them can also overlap, these tracks cannot be treated independently. Instead, for the background model we derive an approach to compute the covariance between tracks analytically and then use this covariance matrix to estimate a combined score (Fig. 1c, Methods S4). We find that even when considering just a single track, our analytical formulation leads to >*7*× speedup over *each* empirical permutation (see Methods S8 and Fig. S1). In practice, numerous shuffles are necessary to compute the mean and variance for a single track, and empirical calculations to estimate the covariance across all tracks is prohibitively slow, highlighting the power and necessity of our analytical formulation.

Though any type of annotated functional region can be incorporated into our framework, here we consider three specific types of tracks. First, **interaction tracks** model various protein–ligand interaction interfaces, where higher positional weights indicate that those positions are more likely to participate in interactions with a ligand; each interaction track corresponds to the subset of protein positions where we have any knowledge about ligand binding potential (as determined by [30, 33]). Second, **domain tracks** span the length of the protein and simply identify portions of the protein sequence that correspond to the domain of interest; weights are 1 for amino acid positions within the domain and 0 elsewhere. Third, the **conservation track** is also the length of the protein sequence, and the weight of each position measures its conservation across vertebrate homologs; higher weights correspond to positions under more evolutionary constraint. Finally, to determine whether a gene as a whole has more mutations than expected, we extend our framework to incorporate the **natural variation track**, which has a single entry per gene that reflects its background mutation rate, as estimated from the number of variants this gene has across healthy populations [35, 36]. Approximately 63% and 90% of human genes have per-site information about interactions or domains respectively, while all genes have per-site conservation values and background gene-level mutuation rates. A gene may have numerous interaction and domain tracks (e.g., for different modeled interaction regions and for each of its identified domains), but has only a single conservation and natural variation track.

The final per-gene score output by PertInInt considers whether somatic mutations across samples are enriched in positions with high ligand-binding potential for an interaction track; within domain positions for a domain track; within conserved sites for the conservation track; and within the gene overall.

### PertInInt effectively identifies cancer driver genes via integrating multiple sources of information

We run PertInInt on somatic point mutation data aggregated across 10,037 pan-cancer tumor samples and 33 tumor types from The Cancer Genome Atlas (TCGA) [2]. PertInInt’s analytical formulation enables the simultaneous consideration of multiple types of biological data regarding protein functionality. However, to first uncover to what extent each source of information—per-site interaction, domain, and conservation information as well overall gene mutational frequency—is independently useful for identifying cancer-relevant genes, we run PertInInt on the pan-cancer dataset when restricted to each of these track types in turn. To validate the method in the absence of a complete gold standard, as we consider an increasing number of output genes, we compute how enriched this set is in genes from the Cancer Gene Census (CGC), a curated list of genes implicated in cancer [37].

We find that utilizing subsets of interaction, domain, conservation, or natural variation tracks can recapitulate known CGC genes to varying degrees, with interaction tracks identifying the largest number of known driver genes while maintaining perfect precision relative to other track subsets (Fig. 2a). Notably, our integrative framework that incorporates all track types outperforms every version of our algorithm that considers only subsets of information; indeed, considering any two sources of biological information outperforms versions of PertInInt that utilize only one source, and considering any three sources of data tends to improve performance even further (Fig. 2b). This demonstrates the ability of our approach to effectively leverage the distinct contributions of multiple, complementary data sources regarding protein position and whole gene functionality in order to uncover cancer driver genes. We note that enrichment of cancer genes amongst PertInInt’s top predictions remains when considering different gold standards (Fig. S3).

**Figure 2:**
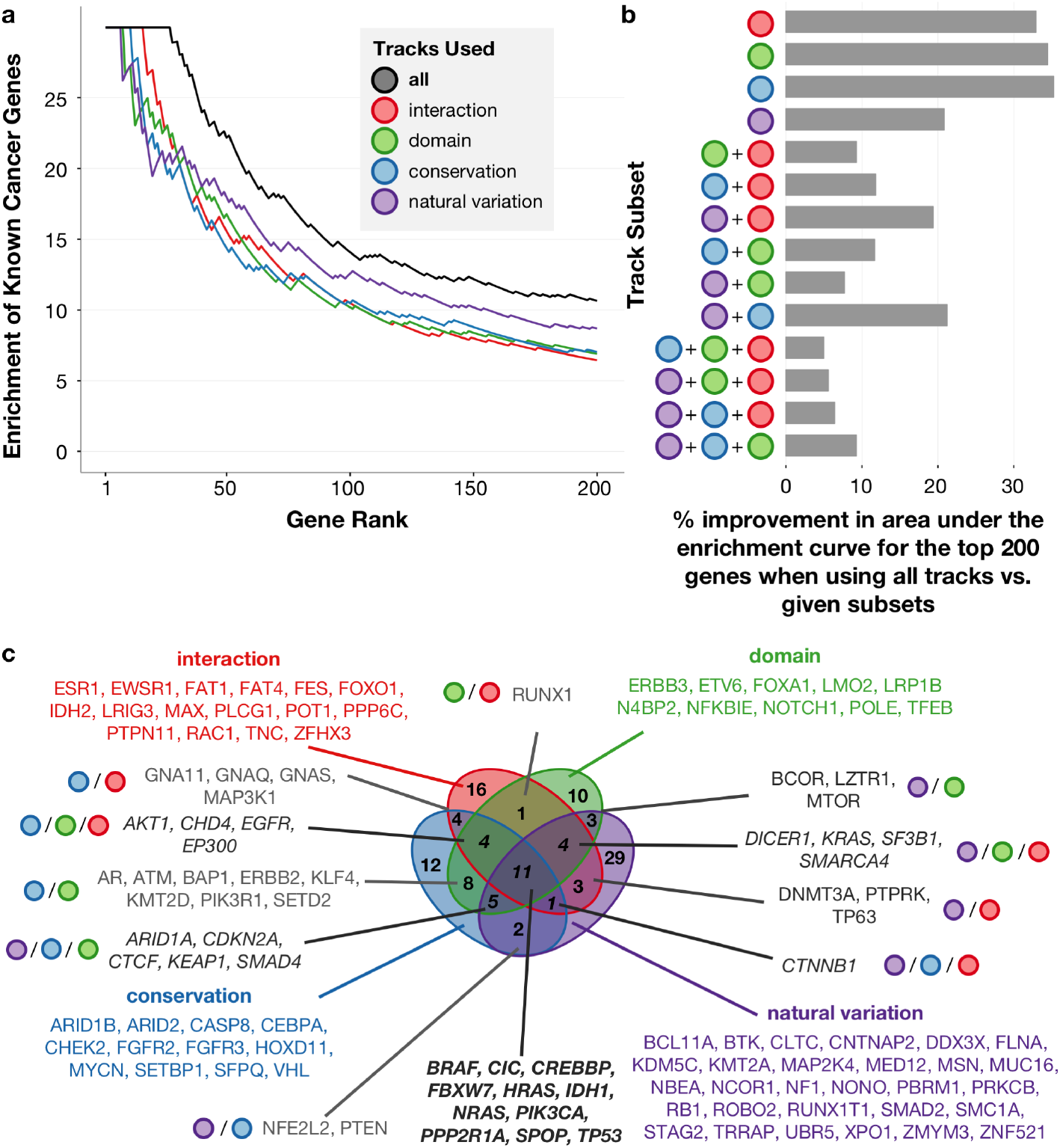
PertInInt is highly effective in uncovering cancer driver genes due to combining multiple sources of information. **(a)** Enrichment of Cancer Gene Census (CGC) genes (*y*-axis) within a given number of top scoring genes (*x*-axis) when run on the pan-cancer dataset using all tracks together (black), only interaction tracks (red), only domain tracks (green), only the conservation track (blue) and only the natural variation track (purple). Enrichment is computed as the ratio between the fraction of CGC genes in the set of top scoring genes considered and the fraction of CGC genes in the whole set of genes. While uncovering genes enriched for somatic mutations within interaction sites, domain positions, conserved sites, or over their lengths each yields cancer-relevant genes, performance is highest when PertInInt uses all sources of information together. **(b)** Percent improvement in the area under the enrichment curve for the top 200 genes when using all track types versus specific subsets of tracks. PertInInt is more effective in uncovering CGC genes when using all sources of information together than when using any other of the possible subsets of information. **(c)** Venn diagram showing the overlap of CGC genes detected in the top 200 genes ranked when considering only interaction, only domain, only conservation, or only natural variation tracks. The different sources of information yield distinct yet overlapping sets of cancer genes.

Strikingly, we find low overlap between the sets of CGC genes identified when utilizing distinct track types, indicating that mutations within cancer genes tend to target a diverse array of functional elements (Fig. 2c). Only a small minority of CGC genes (less than 10%) are identified by all four track types within the top 200 ranked genes. Mutations falling into known tumor suppressor *PTEN*, for instance, tend to hit evolutionarily conserved protein positions but do not alter known inferred interaction interfaces or domain regions more than expected by chance. In contrast, a small molecule binding pocket in the *IDH2* oncogene is recurrently mutated across cancers, and thus it is readily detected using interaction tracks alone but is less significantly ranked when PertInInt is restricted to other functionality data.

### Lowly-mutated genes harbor mutations that preferentially alter functional sites

We next show that PertInInt’s integrative approach can highlight genes with preferentially altered functional sites that may be lowly mutated overall; such “long tail” driver genes are easily missed by traditional frequency-based driver gene detection approaches. When run on the pan-cancer dataset utilizing all track types, PertInInt ranks highly several such infrequently-mutated genes (Fig. 3a). Of the top 35 genes ranked by PertInInt on the pan-cancer dataset, we find that 20 fall into the “long tail” of genes with a missense mutation rate less than one-twentieth of the maximum observed mutation rate (Fig. 3b). These high scoring long tail genes include novel genes with potential implications in cancer as well as known driver genes that cannot have been identified based solely on their relative mutation frequency (e.g., *KMT2D* and *CIC*, Fig. 3b). Many of these infrequently mutated genes harbor significantly perturbed interaction sites, enabling immediate molecular insights regarding their roles in cancer. For example, among long tail genes that are highly ranked by PertInInt but have not yet been identified as cancer-relevant, several have an enrichment of mutations in their DNA or small molecule interaction sites (e.g., *MGA* and *GRIN2D*, Fig. 3b), in line with previous observations that many cancer driver genes exhibit these types of protein interaction perturbations [38–40].

### Mutations are distributed across interaction interfaces

For each protein with a significantly perturbed interaction interface, we next sought to determine whether mutations are found within a small number of interaction sites or across several interaction sites. We consider all sites within the protein with non-zero interaction track weights, and use the frequency with which somatic mutations occur within each of them to compute a normalized Shannon entropy [41]. Higher entropies correspond to proteins with mutations spread across many interaction sites whereas low entropies correspond to mutational patterns that can be uncovered by methods that look for mutation “hotspots” [42]. As expected, PertInInt highly ranks several oncogenes that have previously been detected by hotspot detection algorithms due to their recurrent mutations in critical interaction positions (e.g., *IDH1*, *BRAF*, *NRAS*) [42]. However, there are also many genes with significantly perturbed interaction interfaces where mutations are spread more widely across their interaction sites (Fig. 3c). Known cancer genes *DICER1*, *SMARCA4*, *CREBBP* and *KMT2D*, for instance, are among the top 35 genes ranked by PertInInt and contain significantly mutated interaction sites (combined score across interaction tracks > 6), each with several interaction sites that together harbor an enriched number of somatic mutations.

**Figure 3:**
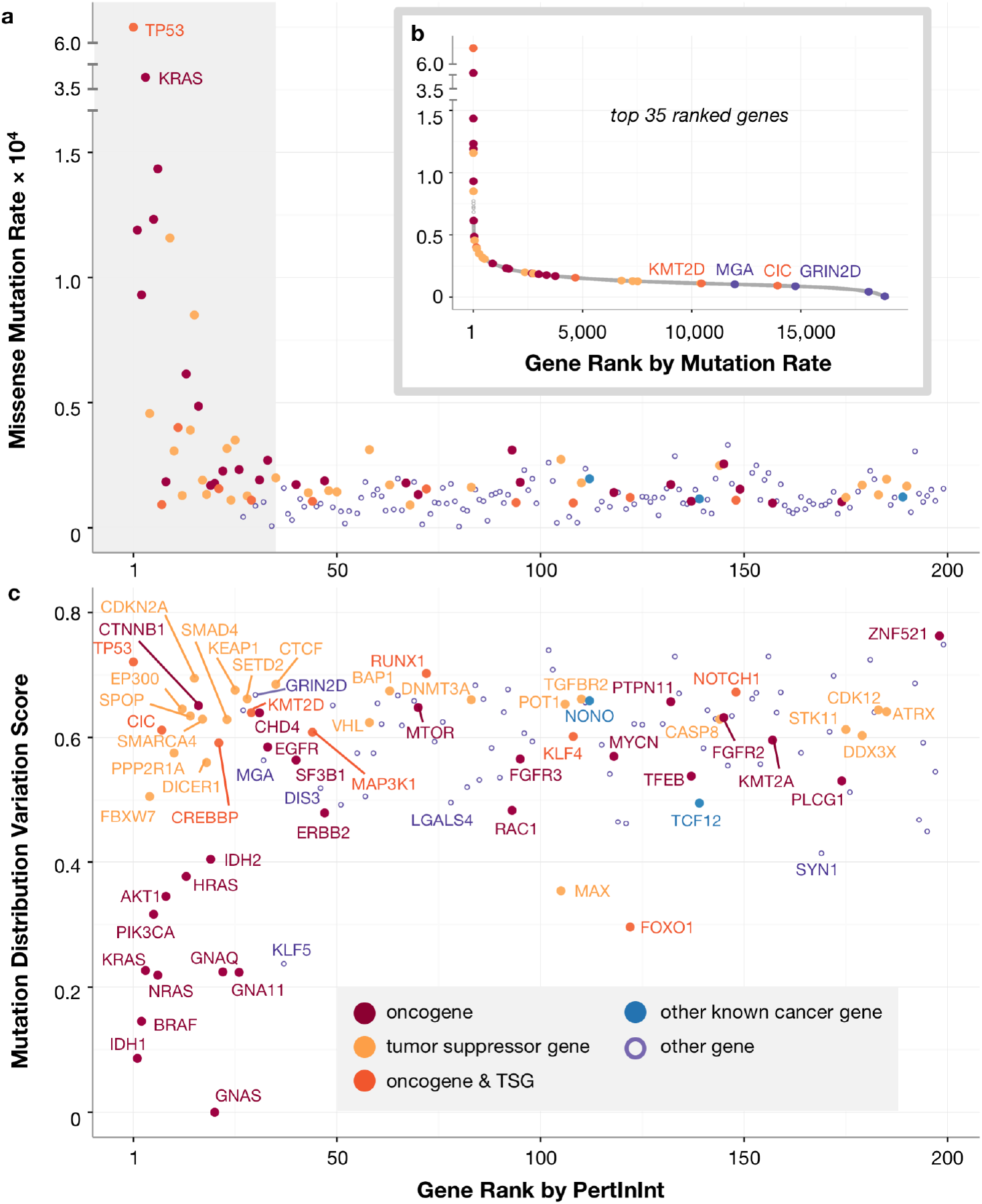
Perturbed interaction interfaces across oncogenes and tumor suppressors. **(a)** Shown are the missense mutation rates (*y*-axis) of the top 200 genes ranked by PertInInt (*x*-axis). Top ranked genes are both highly and infrequently mutated. Genes are colored as in (c). The shaded gray box highlights the plot to 35 genes, which are featured in the part (b) inset. **(b)** Genes are ordered by their missense mutation rate (*x*-axis), and their missense mutation rate is given (*y*-axis). PertInInt’s top 35 ranked genes are plotted in color and exhibit a wide range of ranks with respect to mutation rate. Of these, only genes with below-median overall mutation rates and a *Z*-score ≥ 1 in at least one interaction track are labeled. **(c)** For each of the top 200 genes ranked by PertInInt (*x*-axis), for those with a *Z*-score ≥ 1 in at least one interaction track, we also analyze the distribution of somatic mutations across interaction sites and compute their normalized Shannon Entropy (*y*-axis). These genes contain recurrent (low variation) as well as more distributed (high variation) mutations across their binding interfaces.

Notably, this analysis reveals that the top ranked genes with significantly perturbed interaction interfaces include both oncogenes and tumor suppressor genes (TSGs), reflecting a dichotomy in the impact of binding interface mutations. Whereas some specific mutations within interaction sites have been linked to oncogenic activity [28], other binding site mutations are known to entirely disrupt critical interactions and overall protein function [43]. Although we model the interaction sites of similar numbers of oncogenes and TSGs (238 and 246 respectively), we find that among the 50 genes with the highest enrichment of mutations within their interaction sites, the enrichment of oncogenes is 2.36-fold greater than the enrichment of TSGs. Nevertheless, PertInInt uncovers perturbed interaction interfaces in many genes that have been previously identified as drivers due to nonsense, frameshift, or other relatively disruptive mutations typically associated with TSGs (e.g., *RUNX1* and *FOXO1*). Indeed, enriched yet less common interaction altering missense mutations uncovered by PertInInt may correspond to more subtle knockdown phenotypes or previously underappreciated oncogenic activities of genes traditionally characterized as TSGs.

### PertInInt outperforms previous methods in detecting cancer genes

Having demonstrated that PertInInt can identify interaction interfaces enriched in mutations across tumor samples, and that this is highly predictive of cancer genes, we next turn to assessing PertInInt’s performance as compared to previously published methods for detecting cancer driver genes. These methods differ substantially in terms of their statistical models and overall goals, and the vast majority do not distinguish amongst the various types of interaction and other functional perturbations affecting the identified genes. Nevertheless, we compare PertInInt to other computational methods [15, 16, 21, 35, 42, 44–50] that aim to detect driver genes that either are significantly mutated at the whole gene level, that harbor linear clusters of mutations, that harbor three-dimensional (3D) clusters of mutations, that are enriched for mutations in externally-defined linear regions, or that are enriched for mutations in externally-defined 3D regions (as categorized in [11], see Methods S9).

When applied to tumor samples from the pan-cancer dataset, our method has a greater enrichment for CGC genes than the other tested methods that we were able to run (Fig. 4a). PertInInt also outperforms these methods in terms of enrichment of CGC genes among top ranked genes even after we exclude tumor samples from the six most highly-mutated cancer types with 100+ missense mutations per patient on average, demonstrating that PertInInt’s superior pan-cancer performance is not driven by samples from cancer types that contribute large numbers of mutations (Fig. S4). Notably, the genes ranked highly by PertInInt differ substantially from those identified by other approaches (Fig. 4b). Specifically, the set of genes identified by PertInInt has a consistently low Jaccard Index (JI) with sets of genes ranked by alternate methods (JI < 0.5 across all methods for top 25 genes, JI < 0.25 across all methods for top 150 genes). Importantly, due to our analytical formulation, PertInInt can process the pan-cancer mutational data while considering multiple sources of data about protein functionality in 10 minutes on a single core of standard desktop; alternate methods each consider a limited set of mutational patterns and range in runtime from minutes to days (Table S1).

**Figure 4:**
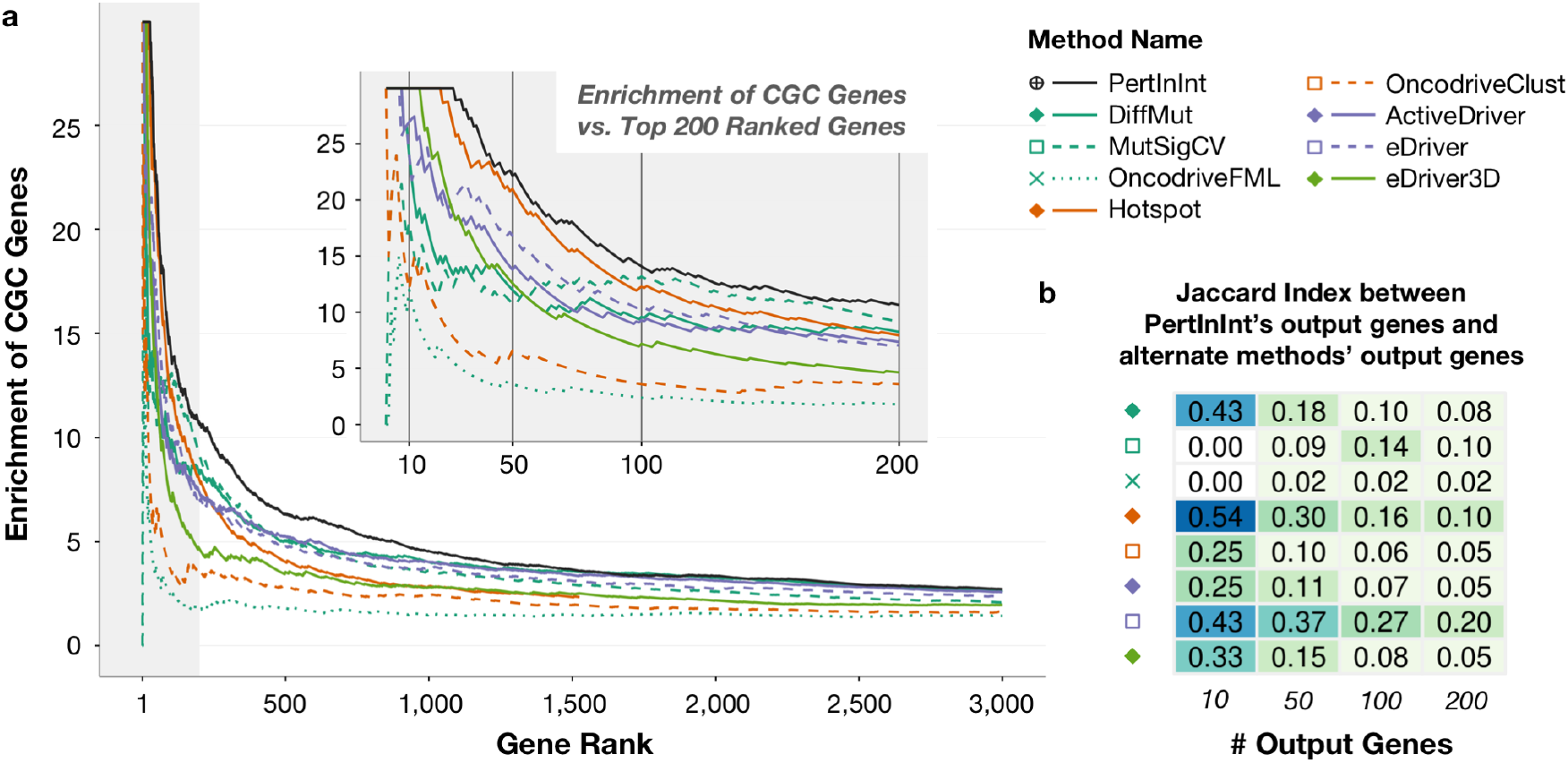
Detection of known cancer genes from a pan-cancer dataset by PertInInt and alternate methods. Each driver gene detection method was run on the pan-cancer set of missense mutations. **(a)** Curves indicate the enrichment of Cancer Gene Census (CGC) genes (*y*-axis) as we consider an increasing number of output genes (*x*-axis) for each driver gene detection method. All methods scored at least 3,000 genes except for Hotspot (orange solid line), which only returned 1,530 genes and whose curve ends at that point. The gray shaded area highlights the plot to 200 genes, a closeup of which is shown in the inset. Vertical lines at 10, 50, 100, and 200 ranked genes in the inset correspond to gene set sizes featured in part (b). **(b)** Jaccard Indices (JIs) are calculated between the top 10, 50, 100, and 200 genes output by PertInInt and the corresponding top 10, 50, 100, and 200 genes output by each other method. Lighter colors indicate lower JIs and less overlap between the gene sets.

We also repeat our analysis on datasets restricted to samples from one cancer type, as many alternate methods that failed to run on the pan-cancer dataset are able to run on these substantially smaller subsets of tumor genomes. We find that in general across individual cancer datasets, PertInInt tends to achieve a higher area under the enrichment curve than other methods, including whole gene methods, and a version of PertInInt that includes only subgene resolution tracks also outperforms other subgene methods (Fig. S5). Overall, these results show that PertInInt is a powerful method for evaluating mutational patterns across tumors of the same cancer type as well as across a pan-cancer dataset covering over 10,000 tumor genomes.

### Distinct perturbed molecular mechanisms uncovered across genes

Having shown that PertInInt is highly effective in identifying cancer genes, we next demonstrate that for each gene highly ranked by PertInInt, we can pinpoint which specific functional regions and mechanisms are perturbed by analyzing each track separately and determining which have positive *Z*-scores. Altogether, we find that that 665 CGC genes have at least one subgene functionality track with a *Z*-score ≥ 0.5, representing functional coverage of 93% of all CGC genes (Fig. 5). Specifically, we find that DNA, RNA, peptide, ion and small molecule interaction sites are enriched in mutations in 16%, 5%, 19%, 14% and 22% of CGC genes respectively; these numbers go up to 23%, 5%, 27% and 24% of CGC genes if including those that are more broadly enriched in mutations across, respectively, DNA-binding, RNA-binding, peptide-binding or metabolite-binding domains (as categorized in Pfam2Go [51]). Up to 77% of CGC genes are enriched in mutations across any domain or interaction site. We note that the perturbed nucleic acid- and small molecule-binding sites or domains found across 45% of cancer genes would not be readily identified by analyses that focus exclusively on protein–protein interaction alterations [16].

**Figure 5:**
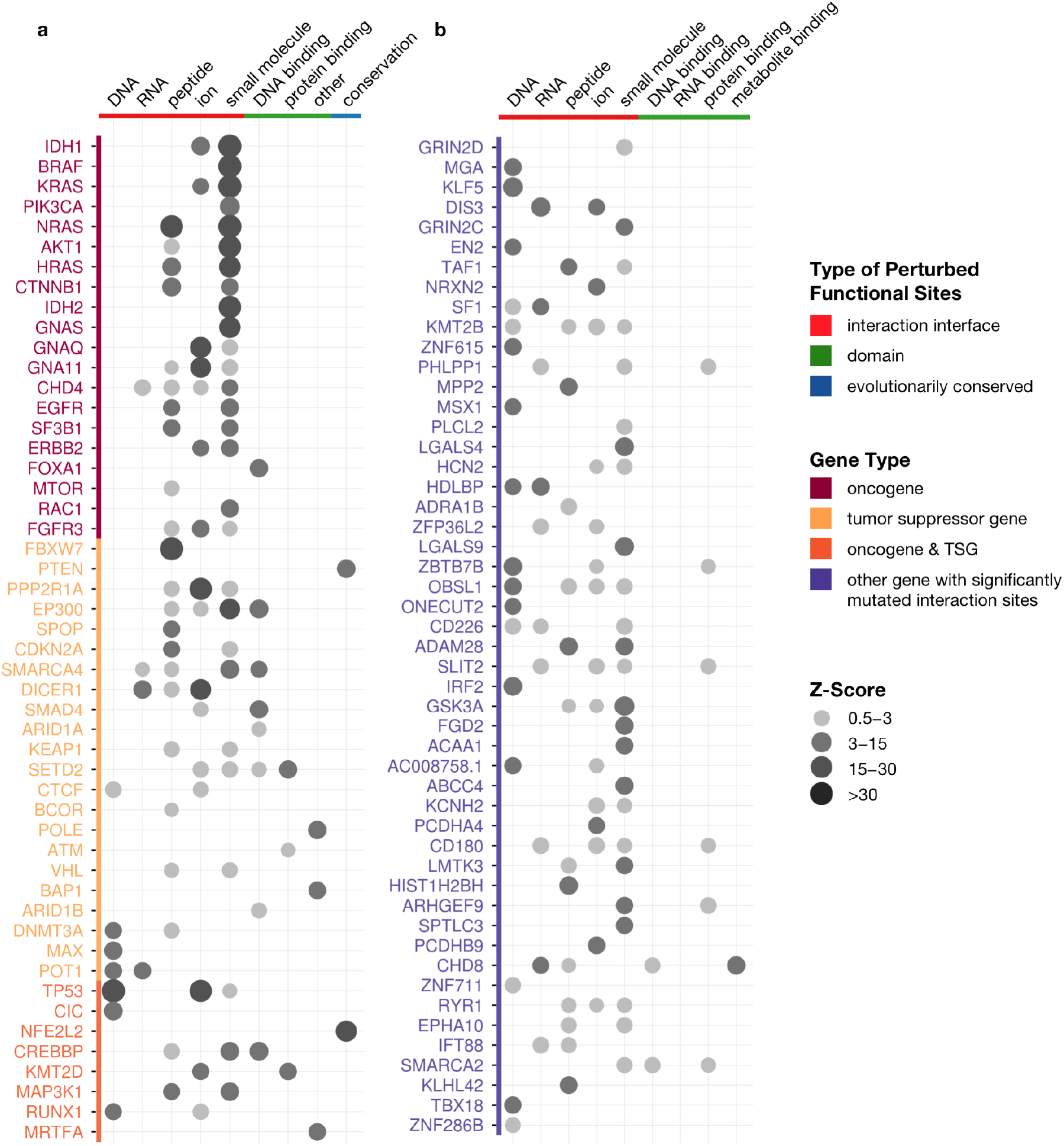
Perturbed mechanisms across oncogenes, tumor suppressor genes, and putative cancer genes. Gene names are colored by driver status; genes that are not yet known to be cancer drivers but have a *Z*-score ≥ 0.5 in one or more interaction tracks are in lavender. For each gene, the circles indicate the *Z*-scores for enrichment of mutations in particular types of tracks, with interaction tracks in red, domain tracks in green and the conservation track in blue. *Z*-scores for mutational enrichments in domain tracks are shown only if the *Z*-scores for the corresponding interaction tracks are < 0.5. *Z*-scores for the conservation track are shown only if *Z*-scores for all other track types are < 0.5. **(a)** PertInInt’s top 50 ranked known cancer driver genes and **(b)** top 50 ranked putative cancer driver genes with a significantly mutated interaction track exhibit a wide range of perturbed functionalities.

We now highlight a few genes that, though not present in the CGC, were uncovered by PertInInt as having significantly mutated interaction interfaces. For instance, transcription factors *MGA* and *KLF5* harbor mutations within their basic helix-loop-helix and C2H2-ZF domains, respectively, that alter their DNA base-binding positions (Fig. 6a), suggesting cancer-specific changes to normal DNA binding and downstream regulatory activity. Indeed, *KLF5*’s E419Q mutation has recently been experimentally shown to change wild-type binding preferences and increase the expression of tumor progression genes *in vivo* [52]. Similarly, *MGA* normally subdues the activity of well-known oncogene *MYC*; its frequent deletion, truncation, or mutated binding properties across cancers further indicates its role as a tumor suppressor [53]. We also find that two RNA-binding genes *DIS3* and *SF1* exhibit significant mutations in their putative RNA-binding sites, with recurrent mutations in *DIS3* altering multiple distinct RNA-contacting positions (Fig. 6b). In support of our predictions, *DIS3* is recurrently mutated in blood and skin cancers and has been identified as a candidate oncogene in colorectal cancer [54]. *SF1* is recurrently mutated across cancers in a mutually exclusive fashion, indicating its analogous functionality, to *RBM10*, a gene found to drive aberrant splicing events in cancer [55].

**Figure 6:**
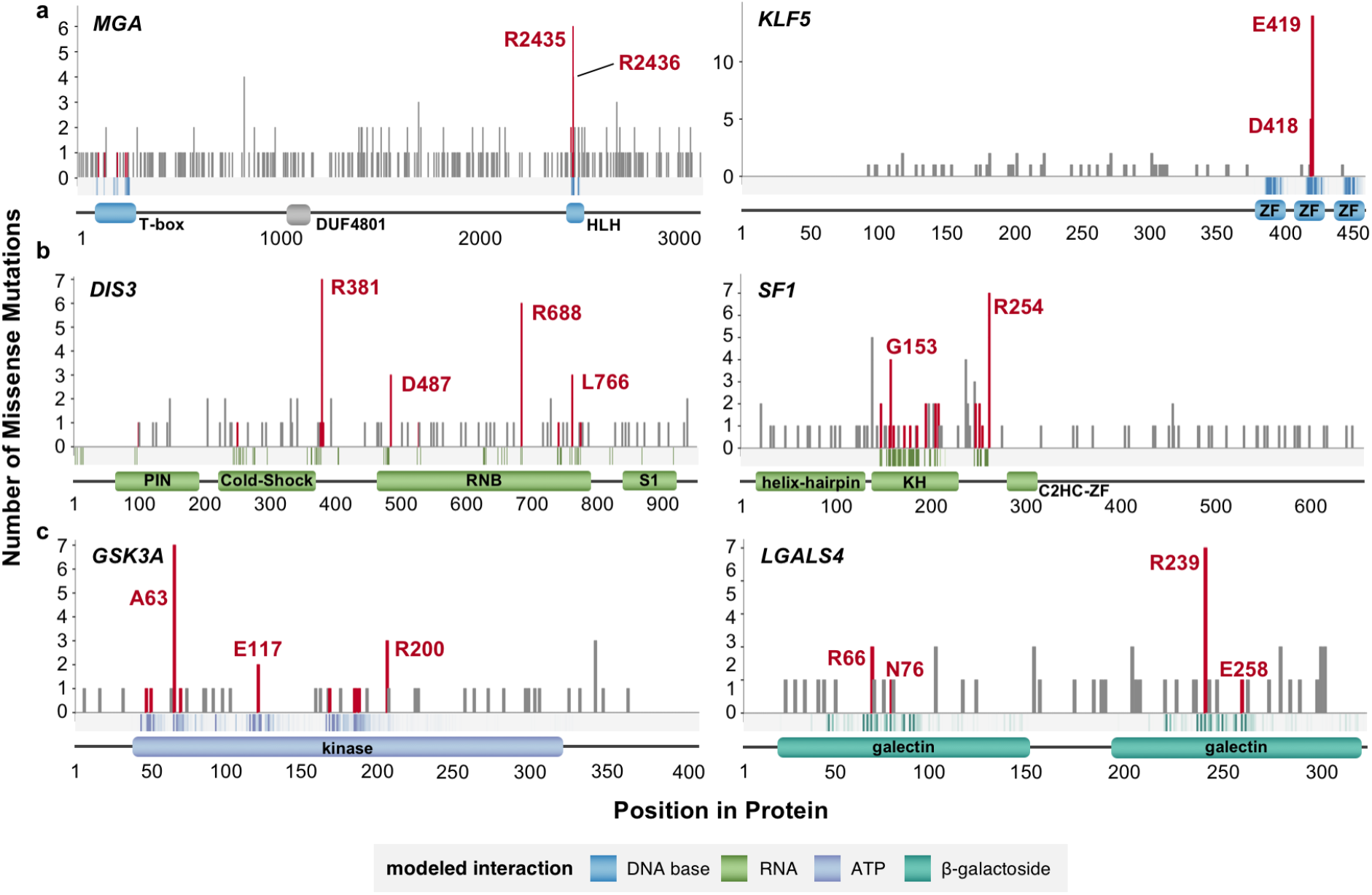
Examples of genes ranked highly by PertInInt that are not known to be drivers. Across the length of each gene (*x*-axis), the number of missense mutations at each protein position is given (*y*-axis). Vertical bars corresponding to mutations affecting binding sites are colored red. The band along the *x*-axis depicts the likelihoods with which residues at each protein position are expected to interact with the specified ligand, with darker bars corresponding to higher (≥0.25) binding likelihoods. Domain locations and names are shown below. **(a)** Putative cancer genes *MGA* and *KLF5* are enriched for mutations in DNA base-binding positions. **(b)** Putative cancer genes *DIS3* and *SF1* are enriched for mutations in RNA-binding positions. **(b)** Putative cancer genes *GSK3A* and *LGALS4* are enriched for mutations in small molecule (ATP and *β*-galactoside sugar, respectively) binding positions.

PertInInt also newly implicates a number of genes—present neither in the CGC nor on other lists of known cancer genes [4, 6, 44, 56]—with mutations that appear to alter critical small molecule binding positions (Fig. 6c). The highly conserved kinase *GSK3A* for instance harbors a significant enrichment of mutations altering its ATP-binding positions. Supporting our prediction, suppression of this gene is associated with impaired growth and induction of apoptosis and it has recently been proposed as a potential therapeutic target in acute myeloid leukemia [57, 58]. We also find that the S-type lectin *LGALS4* has an enrichment of mutations altering the *β*-galactoside sugar-binding positions in its galectin domains; indeed, *LGALS4* has been linked to the regulation of the cancer-relevant Wnt signaling pathway and has been experimentally implicated as a tumor suppressor in colorectal cancer cells *in vitro* [59].

## Discussion

In this work, we have introduced a fast, integrative framework to detect cancer driver genes by uncovering whether somatic mutations across tumors are enriched in sites of different types of functionalities. Our method utilizing this framework, PertInInt, integrates knowledge from the largest set of protein-ligand interaction sites to date [30, 33] with additional biological data regarding subgene functionality and whole gene mutability (Fig. 1). When applied to over 10,000 tumor samples from 33 cancer types, PertInInt reveals a broad range of perturbed functionalities in several known driver genes as well as in relatively rarely mutated genes with predicted tumorigenic roles (Fig. 3a-b, Fig. 5). Notably, PertInInt finds that mutations within many known driver genes are enriched in protein interaction interfaces (Fig. 2a), and more broadly implicates interaction perturbation as a frequent phenomenon in cancer cells (Fig. 5).

Analyses of predicted cancer-relevant coding mutations often involve—whenever possible—assessing their putative effect with respect to protein structure [6, 17, 40, 42, 60]. Although using structure directly to identify relevant mutations is rarely scalable in terms of runtime and coverage [50], PertInInt’s use of structurally predefined regions mediating protein interactions makes large-scale analyses in the context of protein structure feasible. Moreover, since cancer-driving genetic aberrations do not always involve the targeted mutation of protein–ligand interaction interfaces, a critical additional feature of PertInInt—that extends its coverage to all human genes—is that it seamlessly incorporates additional lines of evidence regarding protein site functionality. While here we have demonstrated that PertInInt effectively utilizes per-site evolutionary conservation and domain knowledge, we anticipate that encoding more sources of functional information within our framework (e.g., known phosphorylation sites or intrinsically disordered regions) will unearth other driver mutations and alternate mechanisms of action.

Genes that are frequently mutated across their lengths tend not to overlap genes that exhibit nonrandom patterns of mutations across individual protein positions, a pattern that has previously been leveraged to distinguish tumor suppressor genes from oncogenes [4, 61]. By incorporating whole gene mutability information into our existing framework, we are able to uncover and profile a much more comprehensive set of both oncogenes and tumor suppressor genes (Fig. 2b, Fig. 3c). Although previous methods have also considered the frequency and spatial patterning of mutations within genes together [8, 61, 62], we also simultaneously infer specific perturbed molecular mechanisms within uncovered genes. We note that while mutation deleteriousness predictors—developed both in the context of cancer [25, 26] and otherwise [14]—can evaluate the impact of somatic mutations, they tend to integrate multiple sources of protein site functionality information via complex statistical or machine learning approaches, where the contribution of each data source and thus subsequent mechanistic interpretations are obscured. In contrast, by determining mutational enrichments in specific types of functional sites, PertInInt is able not only to identify cancer-relevant genes but also to begin to explicitly reason about the biomolecular impacts of mutations.

Given the success of large-scale cancer genome sequencing consortia projects in expanding our knowledge of basic cancer biology [63–65], coupled with the decreased cost of genome sequencing, it is clear that sequencing tumor genomes will be routine practice in both basic science and clinical settings, thereby rapidly increasing the number of sequenced tumors available for analysis. Importantly, PertInInt’s analytical framework enables it to efficiently process increasing numbers of tumor genomes; further, this speed is accompanied by better identification of cancer-relevant genes when run on larger numbers of tumor samples (Fig. S6). Since PertInInt’s underlying analytical framework is general, we anticipate that it will also be effective in other settings. For example, because very few non-coding somatic mutations in cancer tend to be recurrent [66], it may be especially powerful for identifying regulatory regions with an enrichment of mutations within sites associated with different measures of functionality (e.g., binding sites for different proteins).

In the future, one of the most tantalizing prospects of cancer genomics is its potential in transforming clinical practice. While identifying and linking cancer mutations to personalized treatments remains a daunting challenge, PertInInt dramatically accelerates the detection of rare mutational driver events from sequenced tumors while providing important information about their mechanisms of action, a key step in developing and customizing targeted therapeutic regimens.

## Supplementary Information

### Supplementary Methods

#### Methods S1. Incorporating per-site information about protein interactions and functionality

Any pre-defined functional region of a protein can be encoded as a track in the PertInInt framework. Currently, we consider three types of per-site functional annotations— interaction, domain and conservation—each of which may yield multiple subgene resolution tracks per protein. Each type of track is described in more detail below.

##### Interaction tracks

Interaction tracks correspond to portions of a protein that are inferred to interact with ligands. These tracks arise in two ways.

First, we utilize the set of “confident” domain–ligand interactions from the InteracDome database (v0.3) [33] to identify putative ligand-binding positions. We use the 9,142 domain–ligand interactions across 1,850 domains with 5+ structural instances. Each position within a domain is associated with a “binding frequency” between 0 and 1 that corresponds to the fraction of the time residues in this position are found in contact with the ligand of interest when analyzing co-crystal structures. For each human protein, we identify instances of InteracDome domains using HMMER (v2.3.2 and v3.1b2), and require complete, high-scoring domain instances as previously described [33, 67, 68]. Within a protein, there is a separate track for each domain–ligand instance within it; this track consists of the residues comprising the match states of the domain, and the weights of these residues are the binding frequencies for the ligand in the corresponding domain positions.

Second, because not all protein interactions are mediated by domains, we leverage sequence homology directly to transfer information from co-complex structures to human protein sequences as previously described [30]. For proteins with one or more regions whose structure in complex with a ligand could be homology-modeled, we introduce a track for each contiguous homology-matched region. Per-position weights reflect the observed residue-to-ligand proximities, computed as the fraction of atoms in the amino acid residue found within 4.0Å of the ligand.

Finally, we note that some domain interactions are mediated not by individual domain instances but by repeating instances of the same domain family. To capture these interfaces, we also consider additional tracks encoding multiple instances of the same domain family in a protein; these tracks span noncontiguous intervals that correspond to the locations of individual domain instances, with track positions weighted according to the binding frequencies at corresponding domain match states as described above. Interaction domain tracks corresponding to domain families with 40+ instances in the same protein are replaced by their aggregate tracks.

##### Domain tracks

For each Pfam-A (v31.0) domain instance within a protein sequence, there is a domain track that specifies which amino acids comprise the domain [67]. Domain tracks span the length of the protein, and positions within and outside of the domain instance are respectively assigned 1- and 0-weights. We again also encode aggregate domain tracks as before to model functional regions mediated by repetitive domain families.

##### Conservation tracks

Each protein has a single conservation track. We obtain the 100-vertebrate cross-species protein multiple sequence alignment from the UCSC Genome Browser [69], and compute per-protein-position conservation-based functionality weights by multiplying the fraction of non-gap residues in the column by the Jensen-Shannon divergence (JSD) between those non-gap residues and a Blosum 62 background amino acid distribution [70].

#### Methods S2. Formulation of per-track somatic mutation functional scores

Suppose we have a protein sequence of length *L* spanning positions *P* = {*p*_1_, …, *p*_*L*_}. This protein is associated with multiple “tracks” *W*, each defined as *W* ⊆ *P*, where each position *p*_*i*_ ∈ *W* is associated with a real-valued weight *w*_*i*_ ∈ [0, 1] reflecting its functionality with respect to the track. Suppose there are *n* cancer somatic missense mutations that fall in positions included in track *W*. For each mutation *i*, let *z*_*i*_ ∈ {*z*_1_, …, *z*_*n*_} be the weight in track *W* of the position where that mutation lies. We further consider the case where each mutation *i* is associated with a value *f*_*i*_ ∈ (0, 1]; here, each *f*_*i*_ is set to the proportion of sequencing reads that contain the mutation (i.e., its subclonal fraction), which has previously been shown to be associated with a mutation’s relevance in cancer [71]. The score of the somatic mutations with respect to track *W* is then defined as:

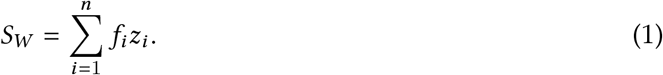

Intuitively, this score reflects the extent to which somatic mutations are falling into functionally important positions within a track.

#### Methods S3. Per-track expectation, variance, and *Z*-score calculation

For a given score *S*_*W*_ for a track, we next want to determine if this score is higher than we would expect by chance. One approach would be to repeatedly randomize the mutations within the positions of the track and use the distribution of resulting scores to compute an empirical *p*-value. Here we show that we can determine the significance of these scores analytically, obviating the need for empirical mutation shuffles and dramatically improving runtime (Fig. S1). Note that in the absence of any selective pressure, the values *z*_1_, …, *z*_*n*_ are independent and identically-distributed (i.i.d.) random variables. We leverage this observation to directly compute the significance of *S*_*W*_. First, we model all mutation locations *z*_*i*_ as being drawn from the same background mutation model *λ*_1_, …, *λ_L_*, where *λ*_*i*_ is the probability that a mutation affects position *i*. If every position *i* within the protein is equally likely to harbor a missense mutation, *λ*_*i*_ = 1/*L*. Here, we incorporate codon-specific missense mutation probabilities as well as cancer-specific C/G-mutation biases into our background mutation model (Methods S10). We linearly scale these values with respect to each track *W* such that 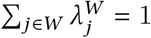. The expected weight of the position in which mutation *i* lies (𝔼[*z*_*i*_]) and its variance 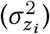 with respect to this null distribution are computed as

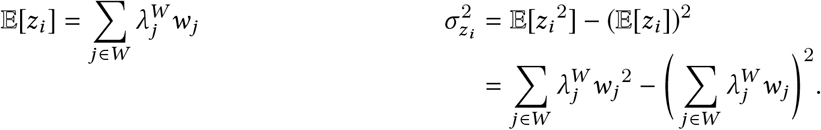

Because the total score of the set of mutations affecting track *W* (i.e., *S*_*W*_) is a sum of independent random variables *f*_*i*_*z*_*i*_ (Eq. 1), the expectation and variance of *S*_*W*_ can also be calculated directly as

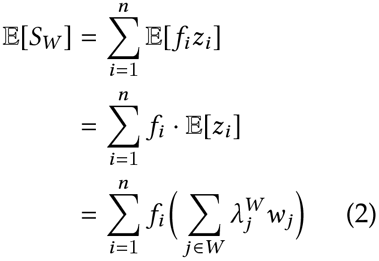

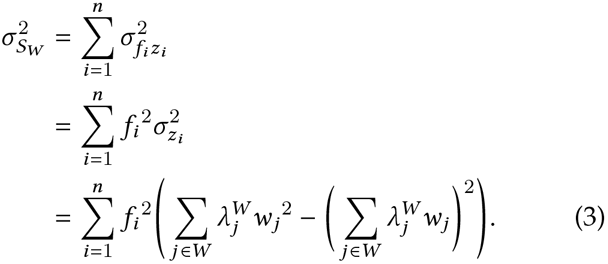

Finally, to determine the significance of the actual score *S*_*W*_, which indicates the propensity of somatic mutations to fall into highly weighted positions in a track, since the sum of independent random variables tends towards a normal distribution, we compute the mutational enrichment *Z*-score for each track *W* as

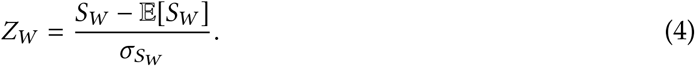

We note that if we restricted each weight within a track to be 0/1 rather than real-valued, restricted mutations to have equal *f*_*i*_ values of 1, and restricted the *λ_i_* to be uniform across the track, we could determine per-track significance analytically using the binomial distribution. Note that with these restrictions, however, we would not be able to incorporate real-valued functionality weights from conservation or interaction tracks, subclonal mutation fractions, or mutational signatures.

#### Methods S4. Between-track covariance calculation

In our framework, a single protein may be associated with *multiple* tracks, each representing a distinct aspect of protein functioning. Since tracks can share positions, the track scores with respect to a set of somatic mutations are not independent of each other, and thus we need to determine their covariance.

Suppose we consider two tracks *V* ⊆ *P* and *W* ⊆ *P*, where each position *p*_*i*_ ∈ *V* is associated with a weight *v*_*i*_ and each position *p*_*i*_ ∈ *W* is associated with a weight *w*_*i*_. Suppose there are *m* mutations (with associated values 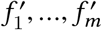) that involve positions within track *V*, and *n* mutations (as before with associated values *f*_1_, …, *f*_*n*_) that involve positions within track *W*. Let *y*_1_, *y*_2_, …, *y*_*m*_ be the weights of the positions that the *m* mutations in track *V* fall into, and let *z*_1_, *z*_2_, …, *z*_*n*_ be the weights of the positions that the *n* mutations in track *W* fall into. Scores are thus calculated as before for tracks *V* and *W* as

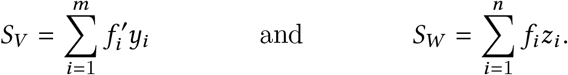

Let *X* = *V* ∩ *W*. If the two tracks do not overlap (i.e., *X* = Ø), then the covariance between *S*_*V*_ and *S*_*W*_ is 0. Otherwise, note that 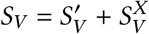, where 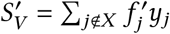 and 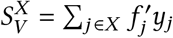. Similarly, 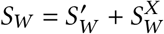. Therefore, we can write covariance as

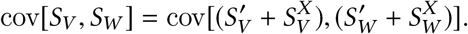

Because the covariance is bilinear, we can now expand this equation as

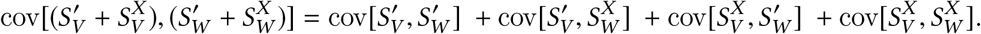

Finally, because mutations landing in track *V* outside of the overlap region *X* have no bearing on *S*_*W*_ and vice versa, the first three covariance terms in the equation above will be evaluated as 0, leaving us with

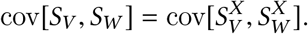

In our framework, we compute covariance conditional on the *q* mutations observed to fall on positions shared by tracks *V* and *W*. Let 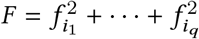, with the *f*_*i*_ associated with the *q* mutations in *X*. With the number of mutations *q* fixed, we have

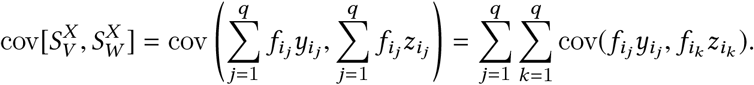

Note that the *same* mutations from tracks 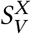 and 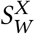 land on the *same* position in the overlap region and simultaneously impact the *S*_*V*_ and *S*_*W*_ scores, whereas any other pair of mutations *j* ≠ *k* are independent. Hence,

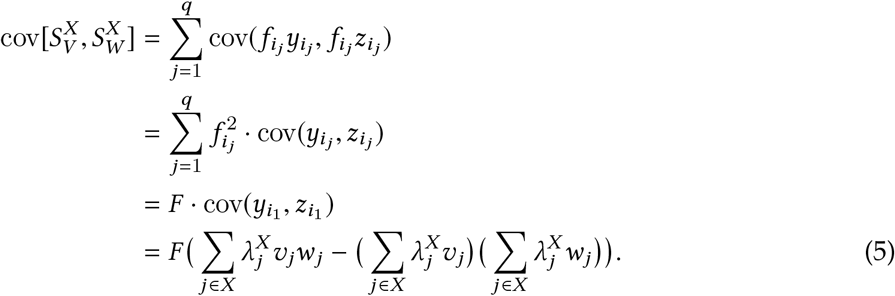

##### Analytical formulation enables precomputation

Remarkably, the per-track expectation, variance and covariance calculations (Eq. 2, 3 and 5) can each be rewritten as *C* · ∑_*i*_ *f*_*i*_ or 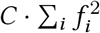, where *C* is fixed per track. We therefore precompute the per-track expectations, variances, and cross-track covariances assuming a single mutation of value 1, and scale these precomputed values at runtime by the mutations observed to fall into each track; this allows PertInInt to achieve an additional 16–18× speedup at runtime (Fig. S7).

#### Methods S5. Incorporating information about overall protein mutability

Using the same analytical formulation described above, we can also compute a *Z*-score per gene reflecting whether the gene is more mutated overall than we might expect. We define a **natural variation track** of length *L* = 19,460^1^ for each gene, where the entry corresponding to the gene of interest has a functionality weight of 1 and all other entries have weights of 0 (i.e., one-hot gene encodings). We then compute a corresponding background mutability probability distribution *λ*_1_, …*λ*_*L*_ based on how much each gene varies naturally across healthy human populations. Specifically, for each of 2,504 individuals included in the 1000 Genomes Project [72], we first min-rank all protein-coding genes by their variant count, linearly scale these ranks to fall between 0 and 1, then round each normalized rank down to its nearest hundredth, which we refer to as its bin. We compute the expected bin value (across individuals) for each gene, and finally to derive the values of *λ*_1_, …, *λ*_*L*_ linearly scale these per-gene expected bin values such that they sum to 1 across all genes. For each track, we use this background mutation model and the *n* mutations observed to fall across all 19,460 genes to analytically compute *per-gene* expectations, variances, and *Z*-scores as before. The covariance between the natural variation track and all subgene tracks is set to 0.

Since the whole-gene track *W* for gene *G*_*j*_ is a one-hot encoding, we can simplify Eq. 1, Eq. 2 and Eq. 3 as

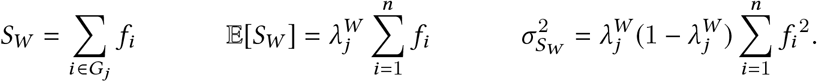

Because the number of mutations affecting all genes is often substantially larger than the number of mutations to affect any single gene, the whole-gene *Z*-scores can be much larger than for the other tracks. We thus effectively subsample the total number of mutations by a factor *s*—set to 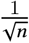 in our implementation—to compute the whole-gene *Z*-scores using the values below before combining them with other subgene *Z*-scores:

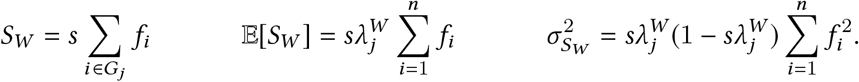

#### Methods S6. Combining multiple per-track *Z*-scores per protein

We evaluate the significance of the scores for all tracks simultaneously using a multivariate normal distribution. Recall that our per-track somatic mutation functional scores (*S*_*W*_, Eq. 1) and their analytically-derived *Z*-scores (Eq. 4) computed for random assignments of mutations are normally distributed when the number of mutations (*n*) is sufficiently large (i.e., by the Central Limit Theorem).

For each track *W*, we empirically determine this minimum *n* by randomly assigning up to 500 mutations to the track 1,000 times in accordance with the corresponding background mutation model (i.e., the *λ*_*i*_’s) and recomputing *S*_*W*_ each time. At each mutation count, we ask whether we can reject the null hypothesis that the mutation functional scores are normally distributed via the Shapiro-Wilk test with *p* < 5e-5. We keep track of the minimum number of mutations per track where we could no longer confidently reject the normality assumption. Only scores derived from mutated tracks with the corresponding required minimum mutation count are modeled together in our multivariate Gaussian. We pre-compute this minimum mutation count value for each track (i.e., before evaluating any cancer somatic mutation data).

For each mutated protein, we compute a single combined score using a weighted *Z*-transform test with correlation correction [73] as

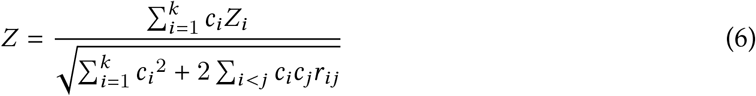

where *k* is the number of tracks with their required minimum mutation count and positive *Z*-scores, *Z*_*i*_ corresponds to the *Z*-score associated with track *i*, *c*_*i*_ is a weight indicating the “confidence” of track *i*, and *r*_*ij*_ is the correlation between tracks *i* and *j* (i.e., *r*_*ij*_ = cov (*S*_*i*_, *S*_*j*_)/*σ*_*i*_*σ*_*j*_). In order to consider each type of functionality data equally, we assign per-track confidences *c*_*i*_ such that the four functional track groups (i.e., interaction, domain, conservation, and natural variation, Fig. 1a) each contribute a quarter of the overall confidence. Within the interaction and domain track groups, where there may be 2+ tracks per group, confidence weights are assigned proportionally to 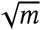, where *m* is the total number of mutations to fall into positively-weighted positions in the track. Finally, we assign a single score per gene by taking the maximum combined score achieved by any of its corresponding protein isoforms.

#### Methods S7. Cancer mutation data preparation

We downloaded all open-access TCGA somatic exome mutation data and RNA-seq expression data from NCI’s Genomic Data Commons on July 15, 2017 [74, 75]. We convert gene expression values (FPKM) to transcripts per million (TPM) and exclude mutations from genes that were expressed at <0.1 TPM in the corresponding tumor sample. For the 765 samples with missing expression data, we exclude mutations from genes that were expressed at <0.1 TPM on average across other tumor samples of the same tissue type. These steps resulted in a filtered set of 1,141,609 missense, 442,070 silent, and 94,813 nonsense mutations across 18,613 genes from 33 cancer types (Fig. S2); note that we consider the unfiltered set of 1,473,729 missense, 578,407 silent and 118,921 nonsense mutations across 19,550 genes when running alternate methods and when running PertInInt to compare to alternate methods. We combine COAD and READ cancer types into the COADREAD group, and GBM and LGG cancer types into the GBMLGG group for per-cancer performance testing (Fig. S5).

#### Methods S8. Runtime Analysis

PertInInt, as well as all algorithm variants of PertInInt and all alternate methods, are run as sole processes on single CPUs, each with a 2.4–2.7 Ghz processor and 30GB of RAM. Methods are timed using Python’s `time` package, and the real (i.e., “wall clock”) elapsed time is reported.

#### Methods S9. Selection and testing of related driver detection methods

We classify alternate cancer driver detection methods based on the mutational patterns they detect; these include whole gene enrichment, *de novo* linear clustering, enrichment in linear externally defined regions, *de novo* three-dimensional (3D) clustering, or enrichment in 3D externally defined regions (as in [11]). We include methods from each of these five groups that require only mutational and/or structural input from the user and have open-source implementations that run locally on a 64-bit Linux machine using sample input. We test the whole gene methods DiffMut [35], MutSigCV [44], and OncodriveFML [45]; the linear clustering methods Hotspot [42], OncodriveClust [46], and NMC [47]; the linear externally defined regions methods ActiveDriver [23], eDriver [21], and LowMACA [48]; the 3D clustering methods GraphPAC [49], iPAC [50], and SpacePAC [15]; and the 3D externally defined regions method eDriver3D [16]. We note that in addition to overall mutation frequency, MutSigCV also considers linear clustering of mutations within genes and the functional impact of mutations based on evolutionary conservation.

All methods including PertInInt are run on the same mutation datasets before our filtering step of limiting to missense mutations from expressed genes. Additional data files required for individual methods are obtained from their most recent online repositories or otherwise from their original publications. For 3D clustering methods, we select a single structural template for each human protein wherever possible as suggested (i.e., preferring native over bound form, longer length, higher sequence identity, higher resolution, and smaller R-value). We note that because these 3D clustering methods only run on proteins with corresponding structural information, their results may be biased toward known cancer genes (i.e., 55.3% of cancer genes have structural templates whereas 28.1% of all genes have structural templates). Methods are run with default parameters, except GraphPAC and SpacePAC, where the significance threshold (*α*) is set to 1.0 to maximize the number of scored genes returned.

For each method, enrichment for CGC genes on increasingly larger sets of predictions is computed; enrichment is computed at each gene rank on pan-cancer results and at every tenth gene on per-cancer results to reduce the impact of minor reorderings of the relatively small number of CGC genes detected across these datasets. Specifically, we calculate enrichment as the fraction of CGC genes in the gene set divided by the fraction of CGC genes in the gene set with 1+ missense mutations; unmutated CGC genes with respect to each mutation dataset are excluded entirely.

Note that LowMACA, NMC, and all three 3D clustering methods did not finish running without error within 30 days on the pan-cancer dataset. We were also unable to run and obtain results from NMC on an additional four individual cancer types (UCEC, SKCM, COADREAD, and LUSC), and thus exclude this method from our evaluation.

#### Methods S10. Background mutational model

We model the likelihoods of protein positions *p*_1_, …, *p*_*L*_ to harbor a missense mutation as *λ*_1_, …, *λ*_*L*_ such that

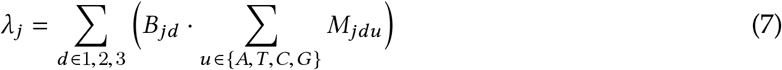

where

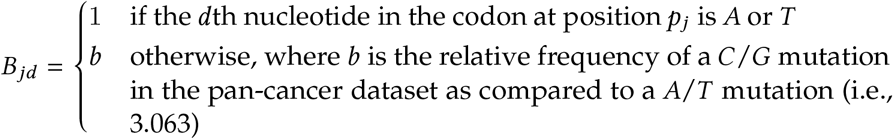

and

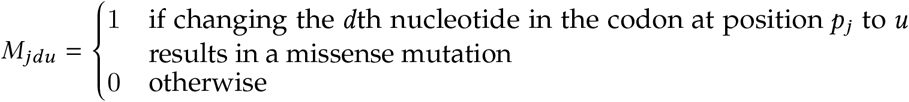

### Supplementary Tables

**Table S1:** Runtimes for cancer driver gene detection methods on pan-cancer dataset. Columns (left to right) are the method classification (as in [11]), method name, and the total time it took for the method to run on the pan-cancer dataset as the sole process on a single node with a 2.4–2.7Ghz processor and 20GB of memory. *MutSigCV failed without 100GB memory allotted. Methods that failed to run without error within 30 days are marked with a ‘--’.

| Method Type | Method Name | Runtime on Pan-Cancer Dataset |
| --- | --- | --- |
|  | PertInInt | 10 minutes, 37 seconds |
| *Whole Gene* | DiffMut | 46 minutes, 51 seconds |
|  | MutSigCV* | 8 hours, 52 minutes, 26 seconds |
|  | OncodriveFML | 32 minutes, 2 seconds |
| *Linear Clustering* | Hotspot | 7 days, 2 hours, 36 minutes, 41 seconds |
|  | NMC | -- |
|  | OncodriveClust | 2 minutes, 30 seconds |
| *Enrichment in Externally Defined Linear Motifs* | ActiveDriver | 7 hours, 38 minutes, 23 seconds |
|  | eDriver | 2 minutes, 23 seconds |
|  | LowMACA | -- |
| *3D Clustering* | GraphPAC | -- |
|  | iPAC | -- |
|  | SpacePAC | -- |
| *Enrichment in Externally Defined 3D Motifs* | eDriver3D | 7 minutes, 25 seconds |

### Supplementary Figures

**Fig. S1:**
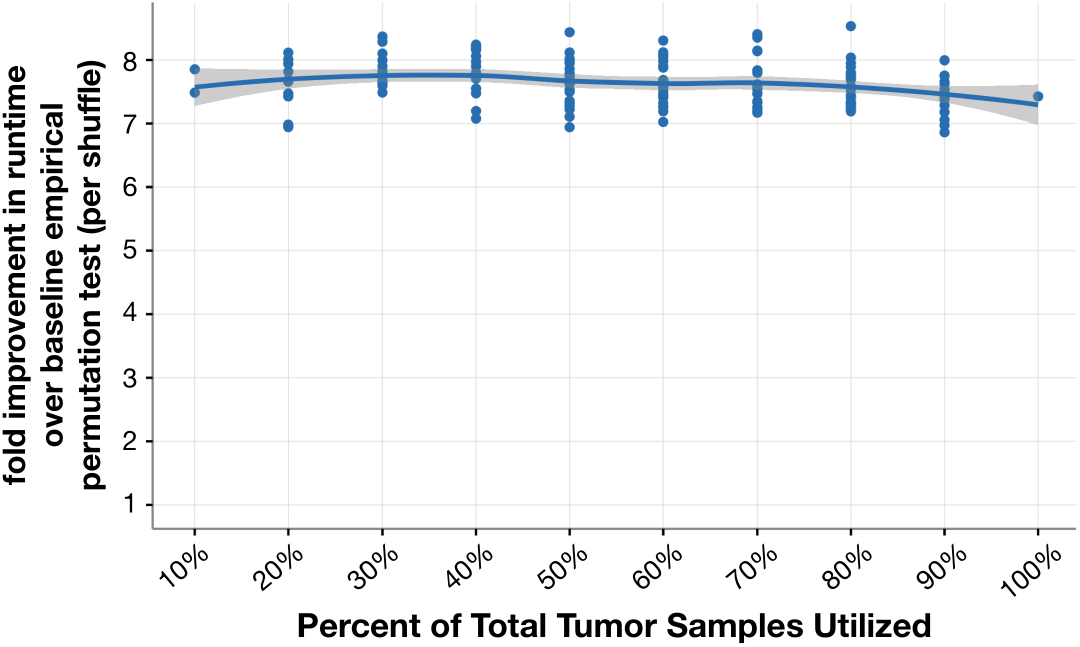
PertInInt’s analytical approach results in >7× speedup over the baseline empirical permutation approach. As a function of the percent (10–100%) of all tumor samples randomly selected from the pan-cancer dataset (*x*-axis), PertInInt’s runtime is compared to a baseline version that uses 1,000 empirical permutations of mutations to estimate *Z*-scores for each track. Shown on the *y*-axis is the fold speedup in runtime for ten random selections of tumor samples of each size. The speedup shown is *per permutation* (i.e., divided by 1,000—the total number of permutations performed across each track). The solid blue line represents the local polynomial regression line, with the gray shading showing standard error. Due to the relatively large runtime of the empirical shuffling procedure, these runtime comparisons use only a single track per protein, conservation.

**Fig. S2:**
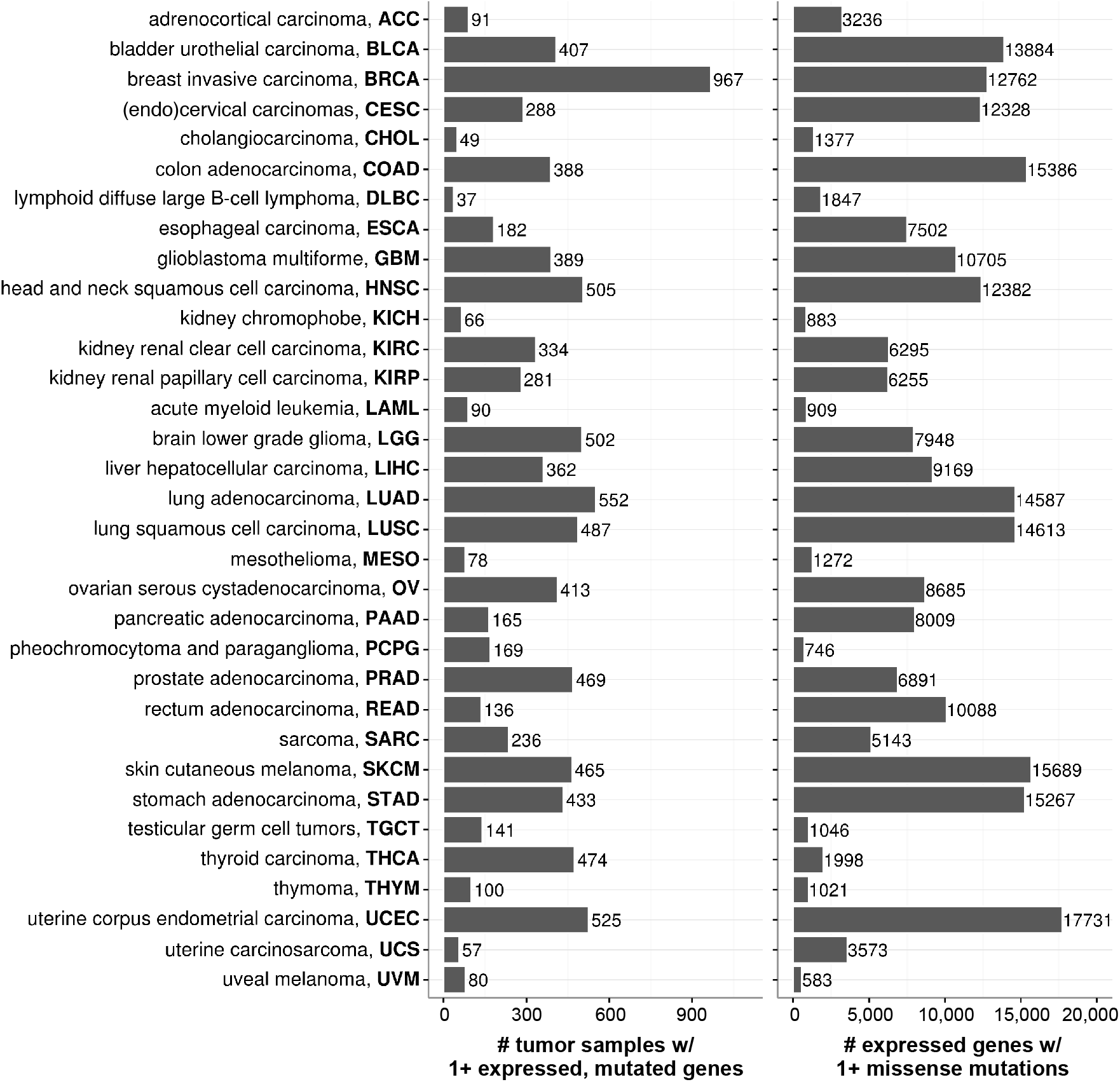
Summary of somatic mutation data. Somatic mutation data obtained from NCI’s Genomic Data Commons Data Portal for 33 cancer types [74]. The number of tumor samples with 1+ expressed (TPM ≥ 0.1) genes with at least one missense mutation is shown in the left plot. The number of genes that are expressed in 1+ tumor samples and have at least one missense mutation is shown in the right plot.

**Fig. S3:**
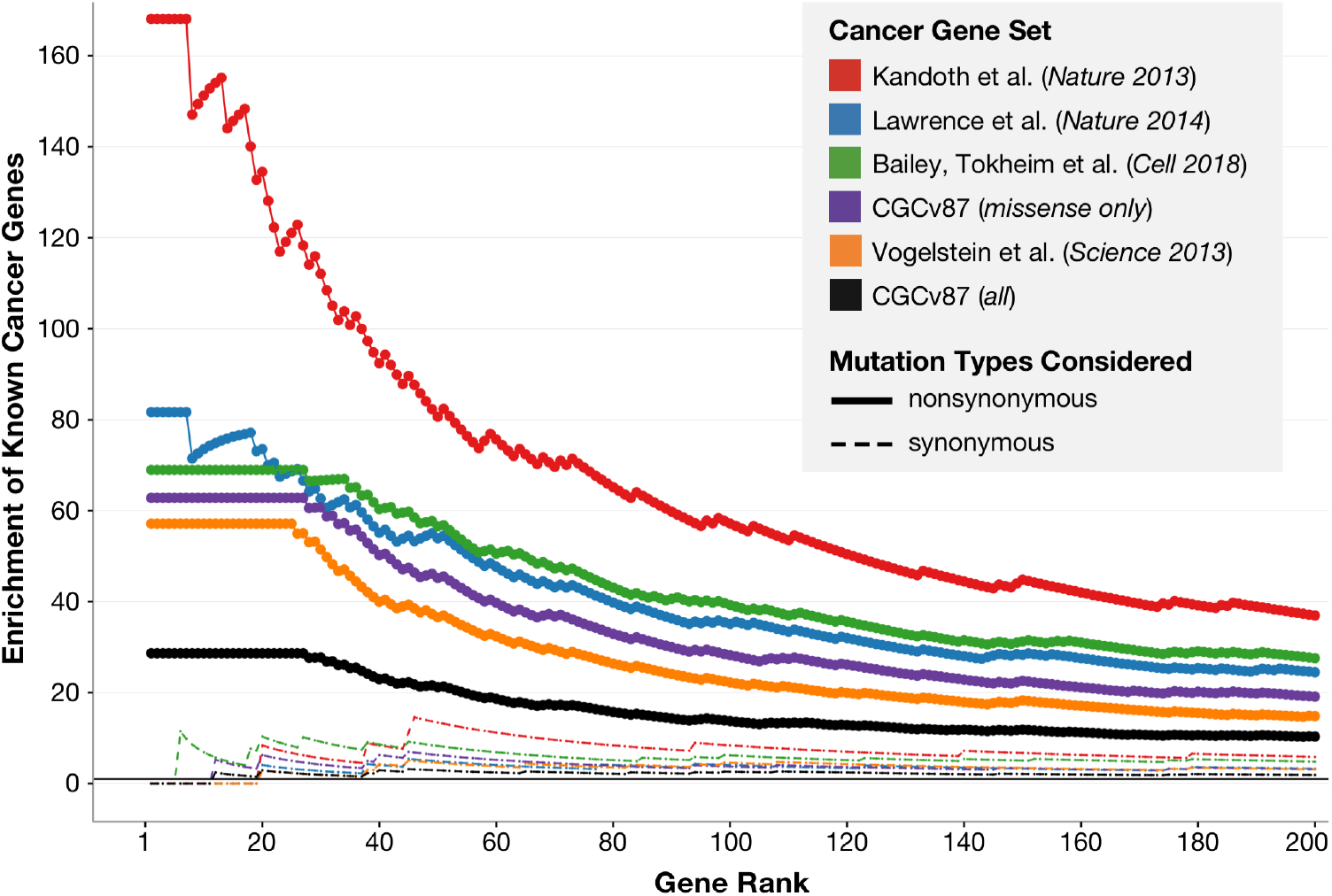
Highly ranked genes are enriched in cancer genes. Gold standard driver gene sets include: those listed in Kandoth et al., 2013 [56] (red), those listed in Lawrence et al., 2014 [44] (blue), those listed in Bailey, Tokheim et al., 2018 [6] (green), all oncogenes and TSGs listed in Vogelstein et al., 2013 [4] (orange), all genes in the CGC (black), and all genes in the CGC with driver statuses due to missense mutations (purple). Ranked gene lists are obtained by applying PertInInt to pan-cancer nonsynonymous mutations (shown as solid lines) and to pan-cancer synonymous mutations (shown as dashed lines). All curves converge to an enrichment of 1 by the end of the ranked list of genes (not shown). Enrichment curves are calculated as described in the main text.

**Fig. S4:**
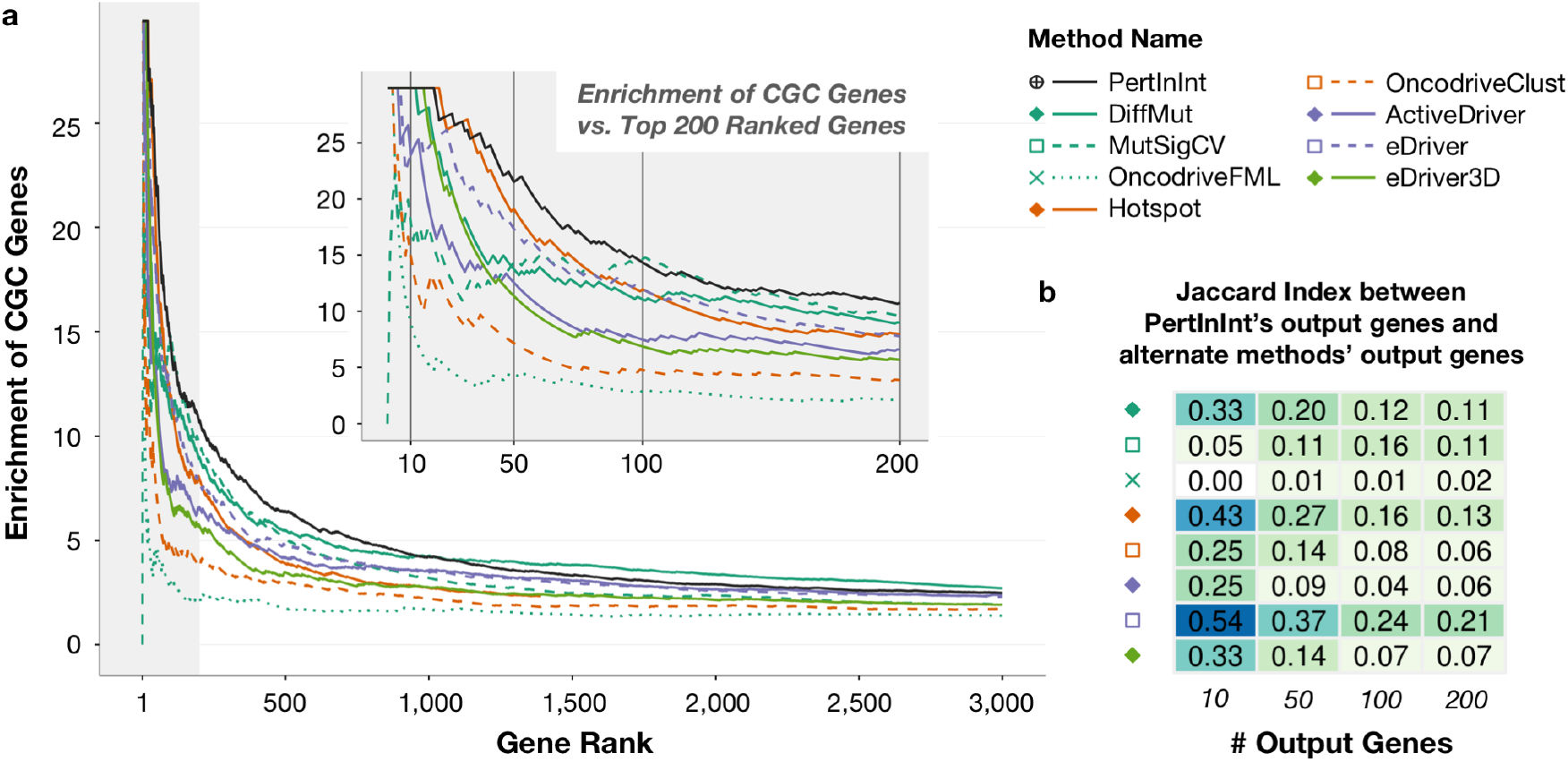
Detection of CGC genes from a pan-cancer dataset excluding highly mutated cancers by PertInInt and alternate methods. Each driver gene detection method was run on the pan-cancer set of missense mutations with tumor samples from highly-mutated BLCA, STAD, SKCM, LUAD, LUSC, and ESCA cancers—where there are more than 100 mutations per tumor sample on average—excluded. **(a)** Curves indicate the enrichment for genes in the Cancer Gene Census (CGC) [37] as we consider an increasing number of output genes for each driver gene detection method. All methods scored at least 3,000 genes except for Hotspot (orange solid line), which only returned 1,397 genes and whose curve ends at that point. The gray shaded area highlights the plot to 200 genes, a closeup of which is shown in the inset. Vertical lines at 10, 50, 100, and 200 ranked genes in the inset correspond to gene set sizes featured in part (b). **(b)** Jaccard Indices (JIs) are calculated between the top 10, 50, 100, and 200 genes output by PertInInt and the corresponding top 10, 50, 100, and 200 genes output by each other method. Lighter colors indicate lower JIs and less overlap between the gene sets.

**Fig. S5:**
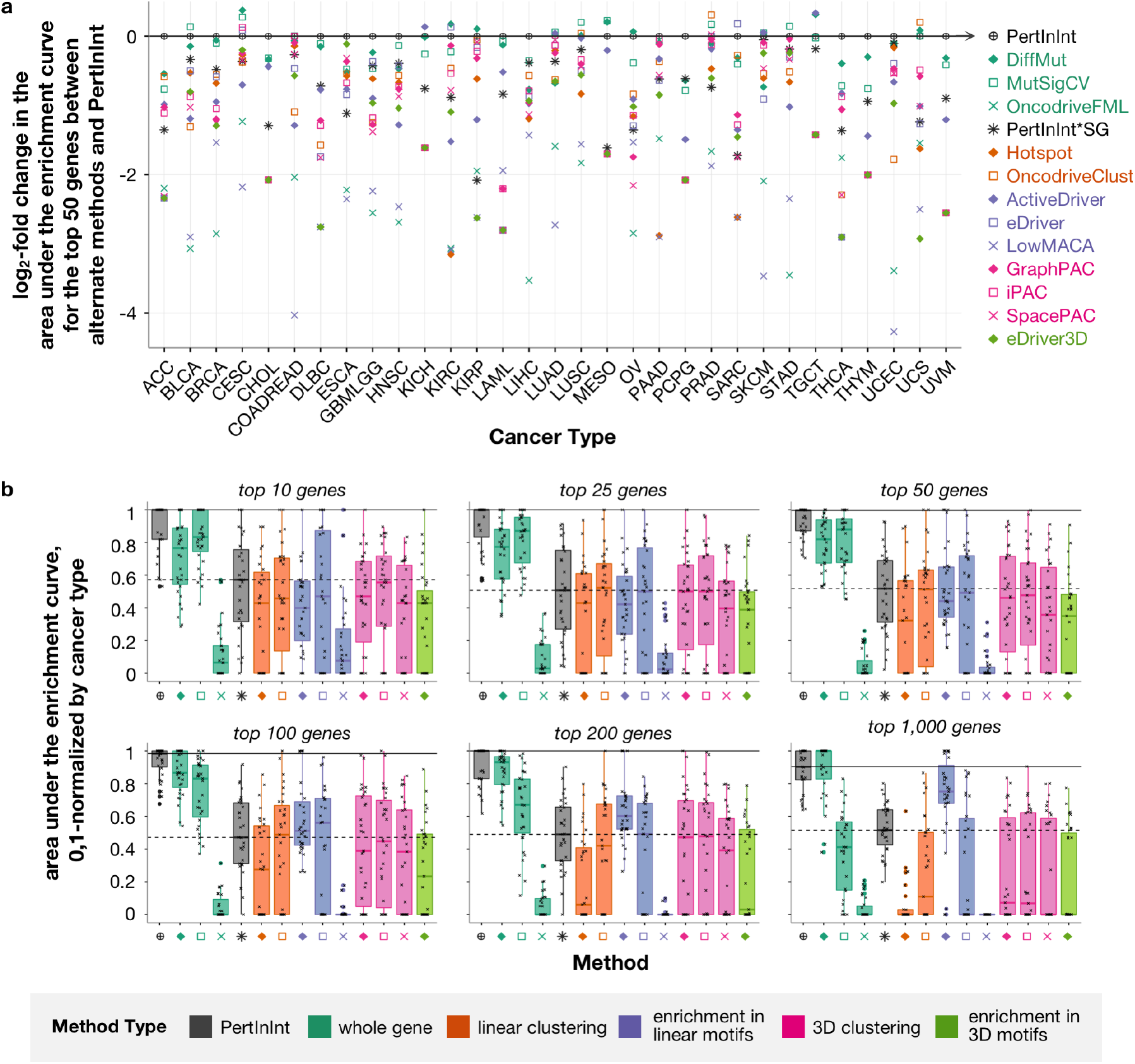
Relative detection of known cancer genes from individual cancer datasets. **(a)** Log_2_-fold change between the area under the enrichment curves for the top 50 genes scored by alternate methods and the top 50 genes scored by PertInInt across individual cancer types. “PertInInt*SG” refers to a version of PertInInt where only subgene resolution tracks are included. PertInInt tends to perform better than the alternate methods, as most of these values are below 0. **(b)** For each cancer type, the areas under the enrichment curves computed for the top 10 (or 25, 50, 100, 200, or 1,000) genes ranked by each driver gene detection method are linearly scaled to fall between 0 and 1. For example, when looking at the top 50 genes ranked by each method when run on SARC mutations, Hotspot has the relatively smallest area under the enrichment curve and thus gets a scaled value of 0, whereas PertInInt has the relatively largest area under the enrichment curve and thus gets a scaled value of 1. Then for each computational method, a box plot of their corresponding values across cancer types is shown. Jittered data points representing different cancer types are overlaid on boxplots. Horizontal solid and dashed lines are drawn at the median relative area under the enrichment curve for PertInInt and PertInInt*SG respectively in each plot. Methods are labeled as in (a).

**Fig. S6:**
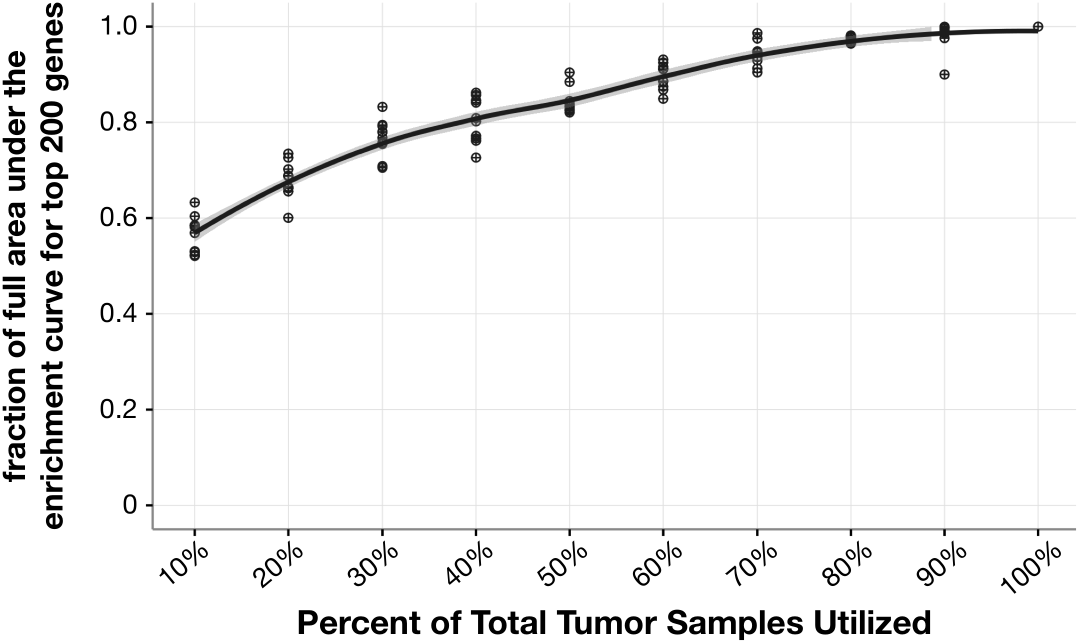
PertInInt’s power increases with more tumor samples. As a function of the percent (10–100%) of all tumor samples randomly selected from the pan-cancer dataset (*x*-axis), we show the area under the enrichment curve for the top 200 genes scored by PertInInt when run on each tumor sample subset, normalized by the area under the enrichment curve for PertInInt’s top 200 predictions when using all tumor samples (*y*-axis). Ten random selections of samples are analyzed at each sample size. The solid black line represents the local polynomial regression line of these normalized areas under the enrichment curve with respect to the sample size. PertInInt’s ability to recapitulate cancer genes increases with sample size.

**Fig. S7:**
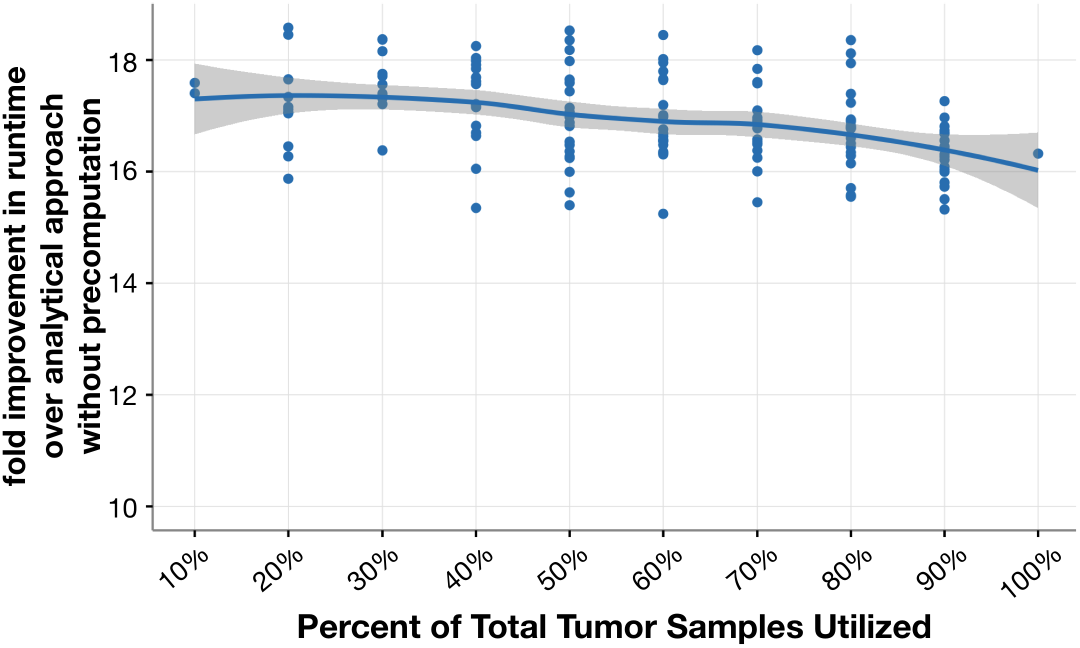
Precomputation enables >16× speedup over basic analytical approach. As a function of the percent (10–100%) of all tumor samples randomly selected from the pan-cancer dataset (*x*-axis), PertInInt’s runtime is compared to a baseline version that does not use precomputed expectation and variance estimates to compute *Z*-scores for each track. Shown on the *y*-axis is the fold speedup in runtime for ten random selections of samples of each size. The solid blue line represents the local polynomial regression line, with the grey shading showing standard error. These runtime comparisons use only a single track per protein, conservation, as in Fig. S1.

## Footnotes

1 PertInInt models 23,278 genes—of which 20,356 are on chromosomes 1–22, X or Y—but only 19,460 genes were profiled in the 1000 Genomes Project, and thus only this many genes have associated natural variation tracks.

## References

[1] International Cancer Genome Consortium, Hudson, T. J., Anderson, W., Aretz, A., Barker, A. D., Bell, C., Bernabé, R. R., Bhan, M. K., Calvo, F., Eerola, I., et al. (2010) International network of cancer genome projects. Nature, 464(7291), 993–998.

[2] TCGA Research Network, Weinstein, J. N., Collisson, E. A., Mills, G. B., Shaw, K. R. M., Ozenberger, B. A., Ellrott, K., Shmulevich, I., Sander, C., and Stuart, J. M. (2013) The Cancer Genome Atlas pan-cancer analysis project. Nat Genet, 45(10), 1113–1120.

[3] Chin, L. and Gray, J. W. (2008) Translating insights from the cancer genome into clinical practice. Nature, 452, 553–563.

[4] Vogelstein, B., Papadopoulos, N., Velculescu, V. E., Zhou, S., Diaz, L. A. J., and Kinzler, K. W. (2013) Cancer genome landscapes. Science, 339(6127), 1546–1558.

[5] McGranahan, N. and Swanton, C. (2017) Clonal Heterogeneity and Tumor Evolution: Past, Present, and the Future. Cell, 168(4), 613–628.

[6] Bailey, M. H., Tokheim, C., Porta-Pardo, E., Sengupta, S., Bertrand, D., Weerasinghe, A., Colaprico, A., Wendl, M. C., Kim, J., Reardon, B., et al. (Apr, 2018) Comprehensive Characterization of Cancer Driver Genes and Mutations. Cell, 173, 371–385.

[7] Garraway, L. A. and Lander, E. S. (2013) Lessons from the cancer genome. Cell, 153(1), 17–37.

[8] Lawrence, M. S., Stojanov, P., Polak, P., Kryukov, G. V., Cibulskis, K., Sivachenko, A., Carter, S. L., Stewart, C., Mermel, C. H., Roberts, S. A., et al. (2013) Mutational heterogeneity in cancer and the search for new cancer-associated genes. Nature, 499(7457), 214–218.

[9] Dees, N. D., Zhang, Q., Kandoth, C., Wendl, M. C., Schierding, W., Koboldt, D. C., Mooney, T. B., Callaway, M. B., Dooling, D., Mardis, E. R., Wilson, R. K., and Ding, L. (2012) MuSiC: identifying mutational significance in cancer genomes. Genome Res, 22(8), 1589–1598.

[10] Torkamani, A. and Schork, N. J. (Mar, 2008) Prediction of Cancer Driver Mutations in Protein Kinases. Cancer Res, 68(6).

[11] Porta-Pardo, E., Kamburov, A., Tamborero, D., Pons, T., Grases, D., Valencia, A., Lopez-Bigas, N., Getz, G., and Godzik, A. (2017) Comparison of algorithms for the detection of cancer drivers at subgene resolution. Nat Meth, 14(8), 782–788.

[12] Reva, B., Antipin, Y., and Sander, C. (2011) Predicting the functional impact of protein mutations: application to cancer genomics. Nucleic Acids Res, 39(17), e118.

[13] Ng, P. C. and Henikoff, S. (2003) SIFT: predicting amino acid changes that affect protein function. Nucleic Acids Res, 31(13), 3812–3814.

[14] Adzhubei, I. A., Schmidt, S., Peshkin, L., Ramensky, V. E., Gerasimova, A., Bork, P., Kondrashov, A. S., and Sunyaev, S. R. (2010) A method and server for predicting damaging missense mutations. Nat Meth, 7(4), 248–249.

[15] Ryslik, G. A., Cheng, Y., Cheung, K.-H., Bjornson, R. D., Zelterman, D., Modis, Y., and Zhao, H. (2014) A spatial simulation approach to account for protein structure when identifying non-random somatic mutations. BMC Bioinformatics, 15(1), 231.

[16] Porta-Pardo, E., Garcia-Alonso, L., Hrabe, T., Dopazo, J., and Godzik, A. (2015) A pan-cancer catalogue of cancer driver protein interaction interfaces. PLoS Comput Biol, 11(10), e1004518.

[17] Kamburov, A., Lawrence, M. S., Polak, P., Leshchiner, I., Lage, K., Golub, T. R., Lander, E. S., and Getz, G. (2015) Comprehensive assessment of cancer missense mutation clustering in protein structures. PNAS, 112(40), E5486–E5495.

[18] Tokheim, C., Bhattacharya, R., Niknafs, N., Gygax, D. M., Kim, R., Ryan, M., Masica, D., and Karchin, R. (2016) Exome-scale discovery of hotspot mutation regions in human cancer using 3D protein structure. Cancer Res, 76(13), 3719–3731.

[19] Niu, B., Scott, A. D., Sengupta, S., Bailey, M. H., Batra, P., Ning, J., Wyczalkowski, M. A., Liang, W.-W., Zhang, Q., McLellan, M. D., Sun, S. Q., Tripathi, P., Lou, C., Ye, K., Mashl, R. J., Wallis, J., Wendl, M. C., Chen, F., and Ding, L. (2016) Protein-structure-guided discovery of functional mutations across 19 cancer types. Nat Genet, 48, 827–837.

[20] Gao, J., Chang, M. T., Johnsen, H. C., Gao, S. P., Sylvester, B. E., Sumer, S. O., Zhang, H., Solit, D. B., Taylor, B. S., Schultz, N., and Sander, C. (2017) 3D clusters of somatic mutations in cancer reveal numerous rare mutations as functional targets. Genome Med, 9(4), 1–13.

[21] Porta-Pardo, E. and Godzik, A. (2014) e-Driver: a novel method to identify protein regions driving cancer. Bioinformatics, 30(21), 3109–3114.

[22] Munro, D., Ghersi, D., and Singh, M. (2018) Two critical positions in zinc finger domains are heavily mutated in three human cancer types. PLOS Computational Biology, 14(6), e1006290.

[23] Reimand, J. and Bader, G. D. (2013) Systematic analysis of somatic mutations in phosphorylation signaling predicts novel cancer drivers. Mol Syst Biol, 9(1), 637.

[24] Zhao, J., Cheng, F., and Zhao, Z. (2017) Tissue-Specific Signaling Networks Rewired by Major Somatic Mutations in Human Cancer Revealed by Proteome-Wide Discovery. Cancer Res, 77(11), 2810–2821.

[25] Shihab, H. A., Gough, J., Cooper, D. N., Day, I. N., and Gaunt, T. R. (2013) Predicting the functional consequences of cancer-associated amino acid substitutions. Bioinformatics, 29(12), 1504–1510.

[26] Carter, H., Chen, S., Isik, L., Tyekucheva, S., Velculescu, V. E., Kinzler, K. W., Vogelstein, B., and Karchin, R. (2009) Cancer-specific high-throughput annotation of somatic mutations: computational prediction of driver missense mutations. Cancer Res, 69(16), 6660–6667.

[27] Kar, G., Gursoy, A., and Keskin, O. (2009) Human cancer protein-protein interaction network: a structural perspective. PLoS Comput Biol, 5(12), e1000601.

[28] Stehr, H., Jang, S. H., Duarte, J. M., Wierling, C., Lehrach, H., Lappe, M., and Lange, B. M. (2011) The structural impact of cancer-associated missense mutations in oncogenes and tumor suppressors. Mol Cancer, 10, 54.

[29] Nishi, H., Tyagi, M., Teng, S., Shoemaker, B. A., Hashimoto, K., Alexov, E., Wuchty, S., and Panchenko, A. R. (2013) Cancer missense mutations alter binding properties of proteins and their interaction networks. PLoS One, 8(6), e66273.

[30] Ghersi, D. and Singh, M. (2014) Interaction-based discovery of functionally important genes in cancers. Nucleic Acids Res, 42(3), e18.

[31] Gress, A., Ramensky, V., Büch, J., Keller, A., and Kalinina, O. V. (2016) StructMAn: annotation of single-nucleotide polymorphisms in the structural context. Nucleic Acids Res, 44(W1), W463–W468.

[32] Engin, H., Kreisberg, J., and Carter, H. (2016) Structure-based analysis reveals cancer missense mutations target protein interaction interfaces. PLoS One, 11(4), e0152929.

[33] Kobren, S. N. and Singh, M. (2019) Systematic domain-based aggregation of protein structures highlights DNA-, RNA-, and other ligand-binding positions. Nucleic Acids Res, 47(2), 582–593.

[34] Forbes, S. A., Bindal, N., Bamford, S., Cole, C., Kok, C. Y., Beare, D., Jia, M., Shepherd, R., Leung, K., Menzies, A., et al. (2010) COSMIC: mining complete cancer genomes in the Catalogue of Somatic Mutations in Cancer. Nucleic acids research, p. gkq929.

[35] Przytycki, P. F. and Singh, M. (2017) Differential analysis between somatic mutation and germline variation profiles reveals cancer-related genes. Genome Med, 9, 79.

[36] Lek, M., Karczewski, K. J., Minikel, E. V., Samocha, K. E., Banks, E., Fennell, T., O’Donnell-Luria, A. H., Ware, J. S., Hill, A. J., Cummings, B. B., et al. (2016) Analysis of protein-coding genetic variation in 60,706 humans. Nature, 536(7616), 285–291.

[37] Futreal, P. A., Coin, L., Marshall, M., Down, T., Hubbard, T., Wooster, R., Rahman, N., and Stratton, M. R. (2004) A census of human cancer genes. Nat Rev Cancer, 4(3), 177–183.

[38] Jeggo, P. A., Pearl, L. H., and Carr, A. M. (2016) DNA repair, genome stability and cancer: a historical perspective. Nat Rev Cancer, 16(1), 35–42.

[39] Delgado, M. D. and Leon, J. (2006) Gene expression regulation and cancer. Clin Transl Oncol, 8(11), 780–787.

[40] Raimondi, F., Singh, G., Betts, M. J., Apic, G., Vukotic, R., Andreone, P., Stein, L., and Russell, R. B. (2017) Insights into cancer severity from biomolecular interaction mechanisms. Sci Rep, 6(34490), 1–9.

[41] Shannon, C. (1948) A mathematical theory of communication. Bell System Technical Journal, The, 27(3), 379–423.

[42] Chang, M. T., Asthana, S., Gao, S. P., Lee, B. H., Chapman, J. S., Kandoth, C., Gao, J., Socci, N. D., Solit, D. B., Olshen, A. B., Schultz, N., and Taylor, B. S. (2016) Identifying recurrent mutations in cancer reveals widespread lineage diversity and mutational specificity. Nat Biotechnol, 34(2), 155–163.

[43] Cho, Y., Gorina, S., Jeffrey, P., and Pavletich, N. (1994) Crystal structure of a p53 tumor suppressor-DNA complex: understanding tumorigenic mutations. Science, 265(5170), 346–355.

[44] Lawrence, M. S., Stojanov, P., Mermel, C. H., Robinson, J. T., Garraway, L. A., Golub, T. R., Meyerson, M., Gabriel, S. B., Lander, E. S., and Getz, G. (2014) Discovery and saturation analysis of cancer genes across 21 tumour types. Nature, 505(7484), 495–501.

[45] Mularoni, L., Sabarinathan, R., Deu-Pons, J., Gonzalez-Perez, A., and López-Bigas, N. (2016) OncodriveFML: a general framework to identify coding and non-coding regions with cancer driver mutations. Genome Biol, 17(1), 128.

[46] Tamborero, D., Gonzalez-Perez, A., and López-Bigas, N. (2013) OncodriveCLUST: exploiting the positional clustering of somatic mutations to identify cancer genes. Bioinformatics, 29(18), 2238–2244.

[47] Ye, J., Pavlicek, A., Lunney, E. A., Rejto, P. A., and Teng, C.-H. (2010) Statistical method on nonrandom clustering with application to somatic mutations in cancer. BMC Bioinformatics, 11, 11.

[48] Melloni, G. E. M., de Pretis, S., Riva, L., Pelizzola, M., Céol, A., Costanza, J., Müller, H., and Zammataro, L. (2016) LowMACA: exploiting protein family analysis for the identification of rare driver mutations in cancer. BMC Bioinformatics, 17(1), 80.

[49] Ryslik, G. A., Cheng, Y., Cheung, K.-H., Modis, Y., and Zhao, H. (2014) A graph theoretic approach to utilizing protein structure to identify non-random somatic mutations. BMC Bioinformatics, 15(1), 86.

[50] Ryslik, G. A., Cheng, Y., Cheung, K.-H., Modis, Y., and Zhao, H. (2013) Utilizing protein structure to identify non-random somatic mutations. BMC Bioinformatics, 14(1), 190.

[51] Mitchell, A., Chang, H. Y., Daugherty, L., Fraser, M., Hunter, S., Lopez, R., McAnulla, C., McMenamin, C., Nuka, G., Pesseat, S., et al. (2015) The InterPro protein families database: the classification resource after 15 years. Nucleic Acids Res, 43(Database issue), D213–221.

[52] Zhang, X., Choi, P. S., Francis, J. M., Gao, G. F., Campbell, J. D., Ramachandran, A., Mitsuishi, Y., Ha, G., Shih, J., Vazquez, F., et al. (2017) Somatic super-enhancer duplications and hotspot mutations lead to oncogenic activation of the KLF5 transcription factor. Cancer Discov, 8, 108–125.

[53] Schaub, F. X., Dhankani, V., Berger, A. C., Trivedi, M., Richardson, A. B., Shaw, R., Zhao, W., Zhang, X., Ventura, A., Liu, Y., et al. (2018) Pan-cancer Alterations of the MYC Oncogene and Its Proximal Network across the Cancer Genome Atlas. Cell Syst, 6(3), 282–300.

[54] de Groen, F. L., Krijgsman, O., Tijssen, M., Vriend, L. E., Ylstra, B., Hooijberg, E., Meijer, G. A., Steenbergen, R. D., and Carvalho, B. (2014) Gene-dosage dependent overexpression at the 13q amplicon identifies DIS3 as candidate oncogene in colorectal cancer progression. Genes Chromosomes Cancer, 53(4), 339–348.

[55] Seiler, M., Peng, S., Agrawal, A. A., Palacino, J., Teng, T., Zhu, P., Smith, P. G., Buonamici, S., Yu, L., Caesar-Johnson, S. J., et al. (2018) Somatic Mutational Landscape of Splicing Factor Genes and Their Functional Consequences across 33 Cancer Types. Cell Rep, 23(1), 282–296.

[56] Kandoth, C., McLellan, M. D., Vandin, F., Ye, K., Niu, B., Lu, C., Xie, M., Zhang, Q., McMichael, J. F., Wyczalkowski, M. A., et al. (2013) Mutational landscape and significance across 12 major cancer types. Nature, 502(7471), 333–339.

[57] Banerji, V., Frumm, S. M., Ross, K. N., Li, L. S., Schinzel, A. C., Hahn, C. K., Kakoza, R. M., Chow, K. T., Ross, L., Alexe, G., et al. (2012) The intersection of genetic and chemical genomic screens identifies GSK-3*α* as a target in human acute myeloid leukemia. J Clin Invest, 122(3), 935–947.

[58] McCubrey, J. A., Steelman, L. S., Bertrand, F. E., Davis, N. M., Sokolosky, M., Abrams, S. L., Montalto, G., D’Assoro, A. B., Libra, M., Nicoletti, F., Maestro, R., Basecke, J., Rakus, D., Gizak, A., Demidenko, Z., Cocco, L., Martelli, A. M., and Cervello, M. (2014) GSK-3 as potential target for therapeutic intervention in cancer. Oncotarget, 5(10), 2881–2911.

[59] Satelli, A., Rao, P. S., Thirumala, S., and Rao, U. S. (2011) Galectin-4 functions as a tumor suppressor of human colorectal cancer. Int J Cancer, 129(4), 799–809.

[60] Niknafs, N., Kim, D., Kim, R. G., Diekhans, M., Ryan, M., Stenson, P. D., Cooper, D. N., and Karchin, R. (2013) MuPIT Interactive: Webserver for mapping variant positions to annotated, interactive 3D structures. Hum Genet, 132(11), 1235–1243.

[61] Tokheim, C. J., Papadopoulos, N., Kinzler, K. W., Vogelstein, B., and Karchin, R. (Nov, 2016) Evaluating the evaluation of cancer driver genes. PNAS, 113(50), 14330–14335.

[62] Korthauer, K. D. and Kendziorski, C. (2015) MADGiC: a model-based approach for identifying driver genes in cancer. Bioinformatics, 31(10), 1526–1535.

[63] Hoadley, K. A., Yau, C., Hinoue, T., Wolf, D. M., Lazar, A. J., Drill, E., Shen, R., Taylor, A. M., Cherniack, A. D., Thorsson, V., et al. (2018) Cell-of-Origin patterns dominate the molecular classification of 10,000 tumors from 33 types of cancer. Cell, 173(2), 291–304.

[64] Ding, L., Bailey, M. H., Porta-Pardo, E., Thorsson, V., Colaprico, A., Bertrand, D., Gibbs, D. L., Weerasinghe, A., Huang, K.-l., Tokheim, C., et al. (2018) Perspective on Oncogenic Processes at the End of the Beginning of Cancer Genomics. Cell, 173(2), 305–320.e10.

[65] Sanchez-Vega, F., Mina, M., Armenia, J., Chatila, W. K., Luna, A., La, K. C., Dimitriadoy, S., Liu, D. L., Kantheti, H. S., Saghafinia, S., et al. (2018) Oncogenic signaling pathways in The Cancer Genome Atlas. Cell, 173(2), 321–337.

[66] Khurana, E., Fu, Y., Chakravarty, D., Demichelis, F., Rubin, M., and Gerstein, M. (2016) Role of non-coding sequence variants in cancer. Nature Review Genetics, 17(2), 93–108.

[67] Finn, R. D., Bateman, A., Clements, J., Coggill, P., Eberhardt, R. Y., Eddy, S. R., Heger, A., Hetherington, K., Holm, L., Mistry, J., Sonnhammer, E. L. L., Tate, J., and Punta, M. (2014) Pfam: the protein families database. Nucleic Acids Res, 42(D1), D222–D230.

[68] Eddy, S. R. (2011) Accelerated profile HMM searches. PLoS Comput Biol, 7(10), e1002195.

[69] Meyer, L. R., Zweig, A. S., Hinrichs, A. S., Karolchik, D., Kuhn, R. M., Wong, M., Sloan, C. A., Rosenbloom, K. R., Roe, G., Rhead, B., et al. (2013) The UCSC Genome Browser database: Extensions and updates 2013. Nucleic Acids Res, 41(D1), 64–69.

[70] Capra, J. A. and Singh, M. (2007) Predicting functionally important residues from sequence conservation. Bioinformatics, 23(15), 1875–1882.

[71] McGranahan, N., Favero, F., de Bruin, E., Birkbak, N., Szallasi, Z., and Swanton, C. (2015) Clonal status of actionable driver events and the timing of mutational processes in cancer evolution. Sci Transl Med, 7(283), 283ra54.

[72] The 1000 Genomes Project Consortium (2012) An integrated map of genetic variation from 1,092 human genomes. Nature, 491(7422), 56–65.

[73] Zaykin, D. V. (2011) Optimally weighted Z-test is a powerful method for combining probabilities in meta-analysis. J Evol Biol, 24(8), 1836–1841.

[74] Fan, Y., Xi, L., Hughes, D. S. T., Zhang, J., Zhang, J., Futreal, P. A., Wheeler, D. A., and Wang, W. (2016) MuSE: accounting for tumor heterogeneity using a sample-specific error model improves sensitivity and specificity in mutation calling from sequencing data. Genome Biol, 17(1), 178.

[75] Grossman, R. L., Heath, A. P., Ferretti, V., Varmus, H. E., Lowy, D. R., Kibbe, W. A., and Staudt, L. M. (2016) Toward a Shared Vision for Cancer Genomic Data. N Engl J Med, 375(12), 1109–1112.

